# Unified somatic calling and machine learning-based classification enhance the discovery of clonal hematopoiesis of indeterminate potential

**DOI:** 10.1101/2024.04.22.590586

**Authors:** Shulan Tian, Garrett Jenkinson, Alejandro Ferrer, Huihuang Yan, Joel A. Morales-Rosado, Kevin L. Wang, Terra L. Lasho, Benjamin B. Yan, Saurabh Baheti, Janet E. Olson, Linda B. Baughn, Wei Ding, Susan L. Slager, Mrinal S. Patnaik, Konstantinos N. Lazaridis, Eric W. Klee

## Abstract

Clonal hematopoiesis (CH) of indeterminate potential (CHIP), driven by somatic mutations in leukemia-associated genes, confers increased risk of hematologic malignancies, cardiovascular disease and all-cause mortality. In blood of healthy individuals, small CH clones can expand over time to reach 2% variant allele frequency (VAF), the current threshold for CHIP. Nevertheless, reliable detection of low-VAF CHIP mutations is challenging, often relying on deep targeted sequencing. Here, we present UNISOM, a streamlined workflow for CHIP detection from whole-genome and whole-exome sequencing data that are underpowered, especially for low VAFs. UNISOM utilizes a meta-caller for variant detection, in couple with machine learning models which classify variants into CHIP, germline and artifact. In whole-exome data, UNISOM recovered nearly 80% of the CHIP mutations identified via deep targeted sequencing in the same cohort. Applied to whole-genome data from Mayo Clinic Biobank, it recapitulated the patterns previously established in much larger cohorts, including the most frequently mutated CHIP genes, predominant mutation types and signatures, as well as strong associations of CHIP with age and smoking status. Notably, 30% of the identified CHIP mutations had <5% VAFs, demonstrating its high sensitivity toward small mutant clones. This workflow is applicable to CHIP screening in population genomic studies.

## INTRODUCTION

Clonal hematopoiesis (CH) of indeterminate potential (CHIP) is a premalignant state, in which hematopoietic stem cells acquire somatic mutations in leukemia-associated driver genes, leading to clonal overrepresentation in peripheral blood. As currently defined, these somatic mutations have variant allele frequencies (VAFs) of 2% or greater, but the individual does not meet the World Health Organization diagnostic criteria for a hematologic neoplasm (1,2). Among the candidate CHIP genes (3), *DNMT3A*, *TET2* and *ASXL1* are most frequently mutated, together harboring over two-thirds of the CHIP mutations identified (4,5). Therein the cutoff of ≥2% VAF for CHIP was chosen due to the limitation of standard next-generation sequencing (NGS) platforms in detecting smaller clones and the presumed rarity of clinical consequences associated with mutations at lower VAFs (5,6).

Biologically, it has been observed that CHIP prevalence increases markedly with age, from <2% of individuals under 50 years of age to approximately 10-20% of those over age 70 (7,8). Importantly, in general population, the presence of CHIP in the peripheral blood of carriers was associated with an 11-fold increased risk of developing hematologic malignancies (7), and two to four times greater risk of cardiovascular diseases (7,9). Furthermore, in non-Hodgkin lymphoma (10) and multiple myeloma patients (11) who received autologous stem-cell transplantation, the prevalence of CHIP at the time of therapy is associated with unfavorable overall survival. Given the biological and medical importance, it is critical to accurately detect CHIP and rare CH.

Accurate screening for CH, including CHIP, requires the detection of low-frequency somatic mutations, which depends heavily on the sequencing platform and depth. Deep targeted sequencing (>400X) can reach lower limits of 0.5-1% VAFs (5,12), and even of 0.2% when combined with single-molecule molecular inversion probes (smMIPs, with >4,000X coverage) (13). Targeted error-corrected sequencing can detect variants with VAFs as low as 0.01% (6), by which at least 95% of the healthy adults above 50 years old were found to carry acute myeloid leukemia (AML) related CH (6,14). Notably, individuals with CH of ≥1% VAF in leukemia driver genes indeed had a significantly increased risk of developing AML (6), which corroborated similar findings in larger AML cohorts (15,16). However, at ∼100X depth of coverage, typically seen in whole-exome sequencing (WES), majority of the CHIP mutations with 2-5% VAFs will not be reliably detected with current somatic calling practice (4), which highlights the need for implementing more robust somatic calling and variant refinement strategies.

Numerous studies have benchmarked the performance of somatic callers over different sequencing depths and VAFs on paired tumor-normal data (17–21). There was obvious discordance among tools, especially at low sequencing depth (17,19,20) and on low-VAF variants (17,18). In fact, a benchmark study by the Pan-Cancer Analysis of Whole Genomes (PCAWG) Consortium revealed that about two-thirds of the discrepancies in mutation calls between WGS and WES from the same cohort are related to low VAFs, and that there is obvious bias from a single caller (22). Thus, ensemble approaches have been proposed to achieve balance between sensitivity and precision (22–24).

Somatic calling in healthy tissues faces additional challenges. Compared to those in tumor samples, somatic variants in normal tissues typically have much lower VAFs due to lower mutation rates (25). However, no analytical pipeline has been developed specifically for CHIP detection and refinement. Instead, a few generic callers, mostly developed for cancer genomes, have been used, such as GATK HaplotypeCaller and UnifiedGenotyper (8), VarScan (26), MuTect together with Indelocator (7,9), and Mutect2 (4,27). Even popular algorithms often lose power on variants with low VAFs (25), including Mutect2 by which majority of the CHIP calls with 2-5% VAFs were not identified at ∼100X sequencing depth (4). Furthermore, refinement of CHIP mutations relying on pre-defined thresholds is less robust and time-consuming.

To systematically address these issues, this study presents **UNI**fied **SO**matic calling and **M**achine learning (ML) based classification, or UNISOM for short, which is a software toolkit designed for streamlined CHIP discovery. UNISOM has implemented a meta-caller to generate an ensemble callset. A ML-based variant classifier was built to separate raw calls into CHIP, germline mutations and artifacts with little manual intervention. When applied to both simulated and real WES/WGS data, UNISOM demonstrated robust performance over a wide spectrum of sequencing depth and VAF.

## MATERIALS AND METHODS

### Leukemia-associated genes and known CHIP

We collected 179 leukemia-associated genes from four CHIP studies with WES (7,8,26) or WGS (4). In addition, we included 189 genes from the Mayo Clinic CHIP diagnostic panel, bringing the total to 202 unique genes (Supplementary Table S1). A list of 2,367 CHIP mutations (1,331 SNVs and 1,036 INDELs) was compiled from three CHIP studies (4,7,8), as described in Supplementary note 1 (“known CHIP mutations”). Over 90% of the CHIP-carriers harbor only a single mutation (4,7,9).

### NA12878 sequencing data

NA12878 is the pilot genome of the Genome in a Bottle (GIAB) Consortium, in which a high-confidence callset has been generated (28,29). This study used four internal datasets (two WES and two WGS), as well as BAMs from five WES (100-360X) and ten WGS (nine with 24-40X and one with 300X coverage) in public sequence repositories (Supplementary Table S2). To reduce computing time and eliminate bias, only alignments falling within the 202 genes were considered. Depending on the original coverage, six WES (>130X) and three (>80X) WGS were downsampled to a coverage series of 200X, 100X, 50X and 20X (Supplementary Table S2). Together, 44 BAMs, 19 from WGS and 25 from WES, were used as input for CHIP spike-in.

### CHIP mutation spike-in

We used BAMSurgeon (v1.2) (30) to add the known CHIP variants to each of the above BAMs, at the same genomic positions where these mutations were originally identified (Supplementary Figure S1). BAMSurgeon used the default parameter settings, except that “the minimum read depth to make mutation” was set at 2 (-m=2) instead of 10 (default). We simulated two batches of BAMs: batch 1 for developing meta-caller and batch 2 for developing ML-based variant classifier (Supplementary Figure S1). Batch 1 started with 12 BAMs from WES and 9 from WGS (Supplementary Table S2). Simulation was performed to either have CHIP-specific VAFs, or have the same VAF each time across all the spike-in CHIP variants. With 13 VAF levels (0.5%, 1% to 10% with 1% increment, 20% and 30%) plus CHIP-specific VAFs, there were 14 simulated data sets varying by VAF, totaling 294 BAMs. Batch 2 was simulated, with CHIP-specific VAFs, from all 44 BAMs (19 WGS and 25 WES). The presence of CHIP spike-in in the output BAMs was verified using bcftools mpileup (31).

### Development of meta-caller

To maximize CHIP detection, we first developed VarTracker based on the idea of force-calling (11,32). It is a permissive tool that uses bcftools mpileup to track mapped reads along genomic positions for the evidence of alternative base(s). VarTracker has five key components: 1) call potential variation sites with bcftools mpileup (“--min-BQ 13”), except in read-dense regions (“-d 1000” by default); 2) use GATK UnifiedGenotyper to genotype candidate variation sites and extract their attributes, such as number of supporting reads, mapping quality, and whether spanning an INDEL site; 3) split multi-nucleotide polymorphisms (MNPs), if any, into individual SNVs; 4) left normalize INDELs; and 5) at multiallelic sites, extract the alternative allele with the most supporting reads, or choose a random one if all alternative alleles have equal number of supporting reads.

To build an ensemble of the most sensitive callers, we benchmarked ten open-source variant callers (Supplementary Table S3), together with VarTracker, on the CHIP genomes from batch 1 simulation. These tools were selected as representatives of widely implemented variant detection algorithms (33). The performance metrics were calculated using the spike-in CHIP as ground truths.

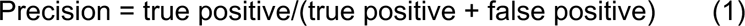

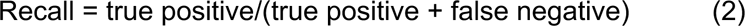

Here, true positive and false negative represent the number of spike-in CHIP that were recovered (at their pre-placed positions regardless of the alternative base) and missed by a caller, respectively, while false positive represents the number of variants called from elsewhere that were not part of the spike-in variant set.

To maximize the recall, a meta-caller was developed by combining VarTracker with Mutect2 and VarDict. These tools showed the highest recall, in particular at the low bound of VAFs (0.5-10%). Of note, Mutect2 and VarDict use haplotype-based strategy (local assembly) and heuristic approach, respectively. VarTracker utilizes bcftools mpileup and GATK UnifiedGenotyper that use Bayesian genotype likelihood model. Thus, the three methods are expected to complement each other in variant detection.

### Development of ML-based variant classifier

To generate variant training sets for benchmarking ML algorithms in variant classification, meta-caller was used to call raw variants from each of the 44 BAMs in batch 2 that were simulated with CHIP-specific VAFs. Raw variants were annotated with Clinical Annotation of Variants (CAVA) (34), and those predicted to have functional effects were retained.

To build prediction models, the retained variants were assigned with an actual class label of being CHIP, GERMLINE, or ARTIFACT, and split into SNVs and INDELs. Variants associated features (26 for SNP and 24 for INDEL) were extracted (Supplementary Table S4). The mlr R package (35) was used to build four ML models, including Recursive Partitioning and Regression Trees (rpart), Support Vector Machine (SVM), Random forest, and eXtreme Gradient Boosting (XGBoost), as well as to perform hyperparameter tuning (k=5, Supplementary Table S5) and calculate feature importance. Three data types (WES, WGS and WES+WGS) were used separately in model building, each with 80% of the raw variants as the training set and the other 20% as the test set. We also tested neural network for CHIP prediction from WGS with TensorFlow platform (https://www.tensorflow.org) in python environment (Supplementary Table S5). Key steps are illustrated in Supplementary Figure S1 and described with more details in Supplementary note 1 (“ML-based variant classifier”).

In calculating the performance metrics, the pre-assigned actual class labels were used to represent the actual class labels. For a given model, its predictive performance was assessed by evaluating the 3-class assignments (i.e., CHIP, GERMLINE, and ARTIFACT) of variants made on the test set against the actual class labels. The prediction outcome was summarized as a 3x3 confusion matrix, which shows the number of raw variants, from a given actual class, that fall into each of the predicted classes. Next, for CHIP, the prediction sensitivity (recall, formula 2), specificity (formula 3), precision (formula 1), F1 score (i.e., harmonic mean of precision and recall, formula 4), and accuracy (formula 5) were calculated based on the confusion matrix.

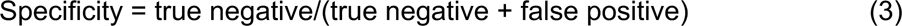

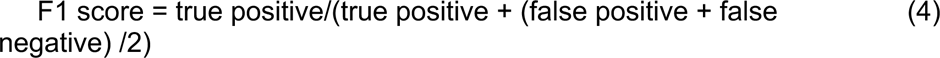

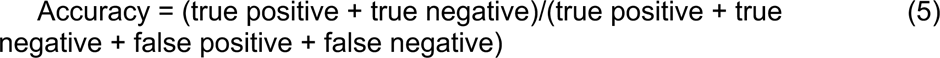

Here, true positive and false negative are the number of actual CHIP that are predicted as CHIP and non-CHIP (i.e., GERMLINE and ARTIFACT), respectively. True negative and false positive represent the number of non-CHIP that are predicted as non-CHIP and CHIP, respectively. For GERMLINE prediction, the metrics were calculated similarly.

### Prediction refinement

XGBoost and Random forest were selected based on the performance metrics. Given the typical coverage of WES/WGS, there is a risk that a small fraction of true CHIP variants might have been erroneously predicted as non-CHIP variants by XGBoost and Random forest. Therefore, we implemented two rules to rescue these predictions. First, non-CHIP variants are flagged as putative CHIP variants if reported in previous studies (4,7,8) and also detected here by at least two of the callers in our workflow. Second, CHIP variants typically have ≤30% VAF (8,36), and variants with higher VAFs are classified as germline by default in our workflow. However, a putative CHIP prediction will be assigned for further review if identified in the five genes (*DNMT3A*, *ASXL1*, *TET2*, *PPM1D* and *JAK2*), as germline variants have rarely been identified in these genes (8).

Following the prediction refinement, a hierarchical Bayesian model was developed to estimate confidence intervals for CHIP predictions (Supplementary note 2). With WES from 75 healthy individuals younger than 40 years of age, the model utilizes CHIP incidence in this reference population as the background (Supplementary note 1, “Confidence interval estimation”).

### XGBoost prediction in WES

The performance of meta-caller and XGBoost classifier was further evaluated on real WES data from a cohort of 25 individuals, for whom targeted deep sequencing data (∼1000X median coverage) were also available. WES had a median coverage of 158X (127-179X). Variants were detected with meta-caller, at default parameter setting, followed by functional annotation with Clinical Annotation of Variants (CAVA) (34). Those predicted to have functional effects were used as input for the XGBoost classifier pretrained on simulated WES data. To assess the prediction, we used the 33 CHIP variants identified from targeted sequencing data as ground truths (Supplementary note 1, “CHIP prediction with WES data”). We used recall, based on formula (2), to measure the predictive performance, where true positive and false negative are those from the 33 mutations that are detected and missed by UNISOM, respectively.

Based on the above analysis of simulated and WES data, we developed the UNISOM pipeline, which includes meta-calling, XGBoost or Random forest prediction, and CHIP prediction refinement and confidence interval estimation (Figure 1A).

**Figure 1.**
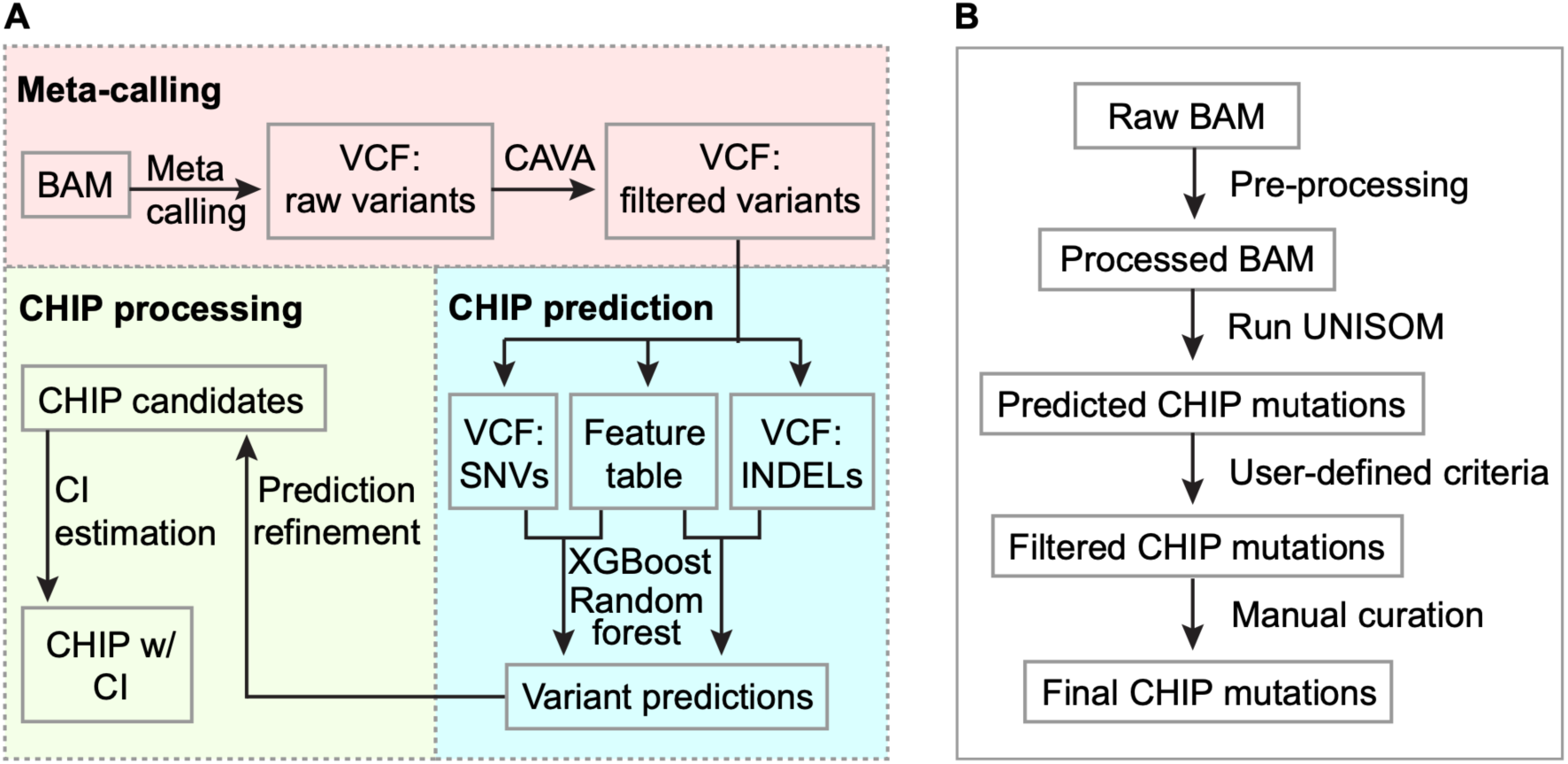
CHIP prediction with UNISOM pipeline. (**A**) Overview of UNISOM pipeline. The pipeline has three major components: meta-calling, ML-based prediction, and CHIP processing. Meta-caller is an ensemble of Mutect2, VarDict and VarTracker. It merges variants from the three tools and assigns a status of 1 to 3 to indicate the number of tools identifying a given variant. Raw variants predicted to have functional effects by CAVA, along with the associated features, are used as the inputs for XGBoost or Random forest prediction. These features, 26 for SNV and 24 for INDEL, cover variant quality metrics, calling status, genomic context, as well as overlap with public variant databases. The prediction model was trained separately for WES versus WGS, and for SNV versus INDEL. The predictions based on XGBoost or random forest model are subjected to further refinement and prioritization. The refinement step recovers those that are most likely to be erroneously predicted as non-CHIP for manual inspection. To enable prioritization of the predicted CHIP, a hierarchical Bayesian model is used to estimate their confidence intervals (CI). The model utilizes CHIP incidence in a population of healthy individuals (here with age <40 years) as the background. (**B**) Key steps recommended for CHIP discovery. To run UNISOM, raw BAM is pre-processed with Picard, which includes coordinates sorting, duplicates marking and filtering based on mapping quality. UNISOM predicted CHIP mutations are then filtered, such as removal of common variants, and extraction of subset based on user provided lists of leukemia driver genes and driver mutations. Finally, to rule out the possibility of being alignment artifacts, it is often necessary to check alignments around the retained variants in a genome browser.

### CHIP prevalence in Mayo Clinic Biobank

To demonstrate its utilization, we applied UNISOM to the ∼30X WGS data from 979 individuals in the Mayo Biobank (37). Library preparation and data processing were reported (38). Prior to CHIP screening, data failed to meet the sequencing quality metrics and the relatedness check, or from participants who self-reported a hematologic malignancy diagnosis in the questionnaire were excluded. UNISOM used default parameters, with the XGBoost classifier trained on WGS data.

Predicted CHIP mutations were removed if: (1) with ≥0.3% MAF in any of the four germline databases (Supplementary Table S4), or with >30% VAF unless identified within the five leukemia driver genes (8) (see “Prediction refinement” section above); (2) low confident variants within low-complexity genomic regions (39); or (3) found in ≥8% of the subjects that likely represent technical artifacts as proposed in (40). The other variants with ≥2% VAF and present in the COSMIC database were retained. To enable comparison with previous cohort studies, we limited our analysis to a list of leukemia driver mutations (4,7), available in Supplementary Table 2 from (4).

## RESULTS

### Overview of UNISOM pipeline

To predict CHIP variants, UNISOM integrates variant calling, annotation, as well as CHIP prediction, refinement and confidence interval estimation in a seamless framework (Figure 1A). It uses BAMs as input and predicts CHIP on a list of leukemia associated genes. The pipeline includes five steps: (1) variants calling with a meta-caller for high sensitivity; (2) variant annotation for quality, genomic context, prevalence in the general population, and impact on protein function using CAVA; (3) annotated variants are classified into CHIP, GERMLINE and ARTIFACT using XGBoost or Random forest based model; (4) rules based refinement to rescue potential CHIP variants from possible mis-predictions (as non-CHIP); and (5) optionally, confidence interval estimation for the predicted CHIP using a hierarchical Bayesian model. We outlined the procedure when applying UNISOM pipeline, from raw alignments to CHIP mutations (Figure 1B).

### Benchmarking of single-sample variant callers on simulated data

To benchmark single-sample variant callers in CHIP detection, particularly at low VAFs (0.5-10%), we simulated a total of 294 CHIP genomes in NA12878 using 13 uniform VAFs, as well as CHIP-specific VAFs by replicating known CHIP mutations. These known mutations have a median VAF of 12.5% from 1,331 SNVs and 14.8% from 1,036 INDELs. By counting the pileup at genomic positions pre-inserted with these known CHIP variants, we checked whether data simulated with CHIP-specific VAFs can capture their actual VAF spectrum. Using sample NA12878_01 as an example (Supplementary Table S2), the SNVs and INDELs had a median VAF of 5.4% and 4.7%, respectively, in the simulated 50X WES data, similar to that in simulated 50X WGS data (4.8% and 6.3%). The trend of lower VAFs, compared to those of the actual spike-in CHIP (Supplementary Figure S2), was observed in all simulated data. This discrepancy is partly due to the fact that not all CHIP mutations have been successfully simulated into the genome, given the low VAFs of many CHIP mutations and insufficient depth of coverage. Of note, the overall lower VAFs in simulated data do not prohibit the benchmarking of callers that have higher sensitivity at low VAFs.

To identify the most sensitive callers, we used the alignments (BAMs) simulated with uniform VAF. Overall, the tools performed comparably in the detection of germline variants; however, marked differences were observed in the detection of somatic mutations. Using the highly confident germline calls (NISTv3.3.2) (29) as the ground truths, all the 11 tools had ≥96% SNV recall rate in 100X WGS and 10/11 tools (except varScan, 92% recall rate) had ≥98% recall rate in 100X WES from NA12878_01. For the detection of spike-in CHIP variants, VarTracker had the highest recall for both SNVs and INDELs, followed by VarDict and Mutect2, over the full range of VAFs tested (Supplementary Figure S3A-D). VarTracker identified 65.7-85.4% of the spike-in SNVs, versus only 27-54.5% by VarDict and Mutect2 (Supplementary Figure S3A and B). A similar pattern was observed for INDELs (Supplementary Figure S3C and D). However, the remaining 8 callers tested had much reduced recall, more obviously at <10% VAFs.

Next, we evaluated variant calling performance with simulated data at different read depths (20-100X) (Figure 2A and B, Supplementary Figure S4A and B). VarTracker, VarDict and Mutect2 had the highest recall for both SNVs and INDELs in WES (Figure 2A) and WGS (Figure 2B) across all depths. Combining WES and WGS for simplicity, VarTracker had a recall of 72.7-92% for SNVs and 92.5-98.8% for INDELs, which was 21-44% higher than that of VarDict and Mutect2. Not surprisingly, of the three callers, Mutect2, which had the lowest recall, had the highest precision, followed by VarDict and VarTracker (Supplementary Figure S4A and B). Previous studies have reported a dependence of calling sensitivity on coverage depth for VarDict (41), Mutect2 (4), as well as Samtools mpileup (i.e., bcftools mpileup) and GATK UnifiedGenotyper (both used by VarTracker) (42). Thus, we also checked how coverage impacts variant calling for these methods. As expected, overall, the number of identified spike-in variants increased with coverage for both WES (Supplementary Figure S5A and B) and WGS (Supplementary Figure S5C and D).

**Figure 2.**
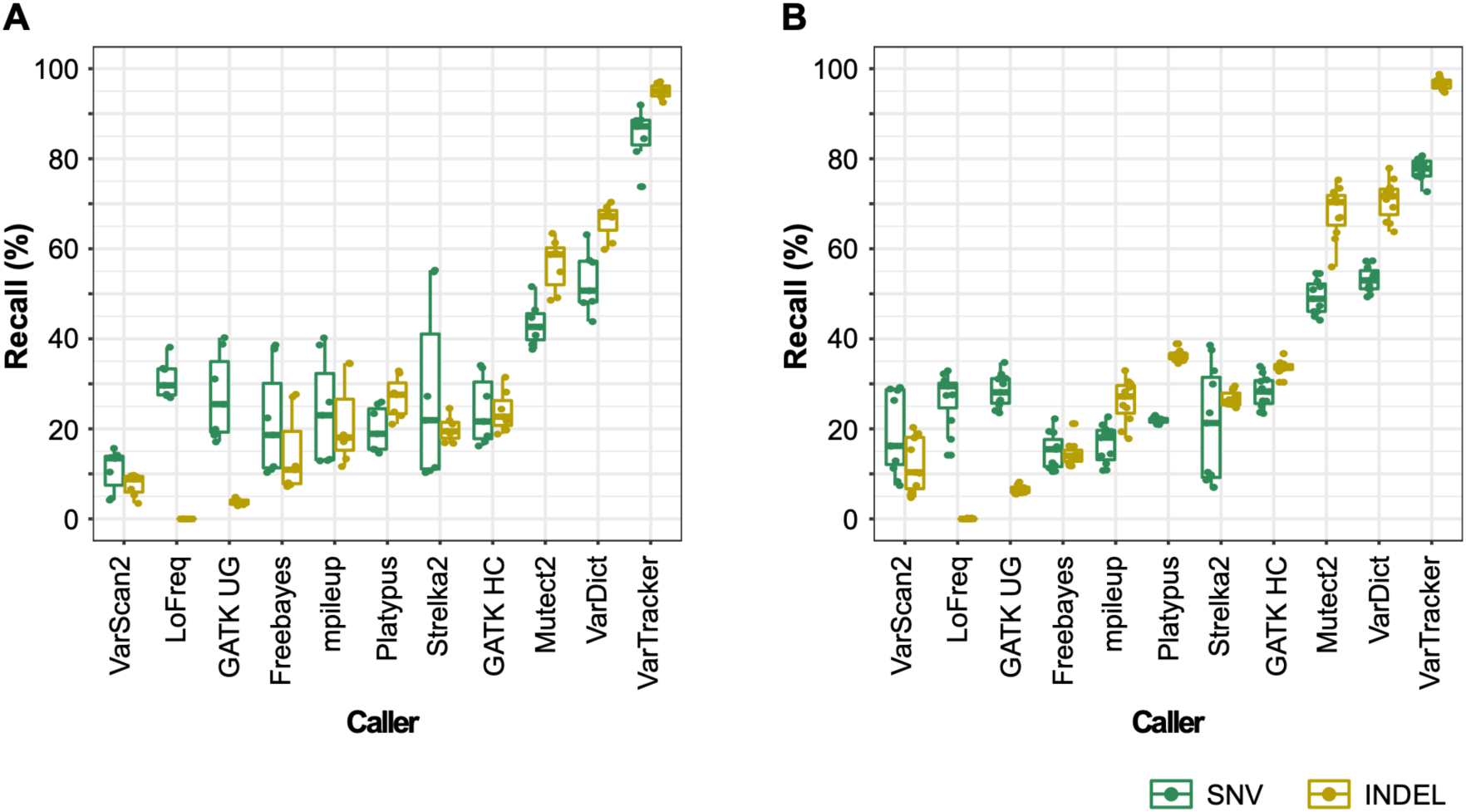
Recall of 11 tools tested on simulation data. (**A**) Simulated SNVs and INDELs in WES. (**B**) Simulated SNVs and INDELs in WGS. Each box plot in (A) and (B) used recall rates estimated from 7 WES and 11 WGS data, respectively, split into SNV and INDEL. All data are from batch 1 simulation that used CHIP-specific VAFs, excluding those with >100X coverage. The 7 WES data are from datasets 13, 18 and 19 with coverage of 20X, 50X and 100X, while the 11 WGS data are from datasets 1, 3, 4, 11, and 12 with coverage of 20X, 29X, 50X, 84X and 100X (See Supplementary Table S2). VarTracker, VarDict and GATK Mutect2 showed the highest recall for both SNVs and INDELs. GATK UG, UnifiedGenotyper; GATK HC, HaplotypeCaller.

The above analyses revealed that VarTracker, VarDict and Mutect2 had the highest recall in the simulated data. Of note, the increased recall from VarTracker is mainly due to the inclusion of calls with a single supporting read, which were not reported by Mutect2 and VarDict. To support this, we checked overlap of the three tools in recovering spike-in from 100X WES (Figure 3A and C) and 50X WGS data (Figure 3B and D) in sample NA12878_02 (Supplementary Table S2). Of the total spike-in, about 90% were recovered by the three tools combined (Figure 3A and B), which dropped to <70% if excludes VarTracker calls with a single supporting read (Figure 3C and D). In fact, about 90% of the VarTracker unique calls (not identified by the other two) had only a single supporting read (Figure 3A and B). At the same cutoff of ≥2 supporting reads, VarTracker and VarDict had comparable recall, which is 3-7% higher than that of Mutect2 (Figure 3C and D). To enhance the variant detection, a meta-caller was built by combining the three tools.

**Figure 3.**
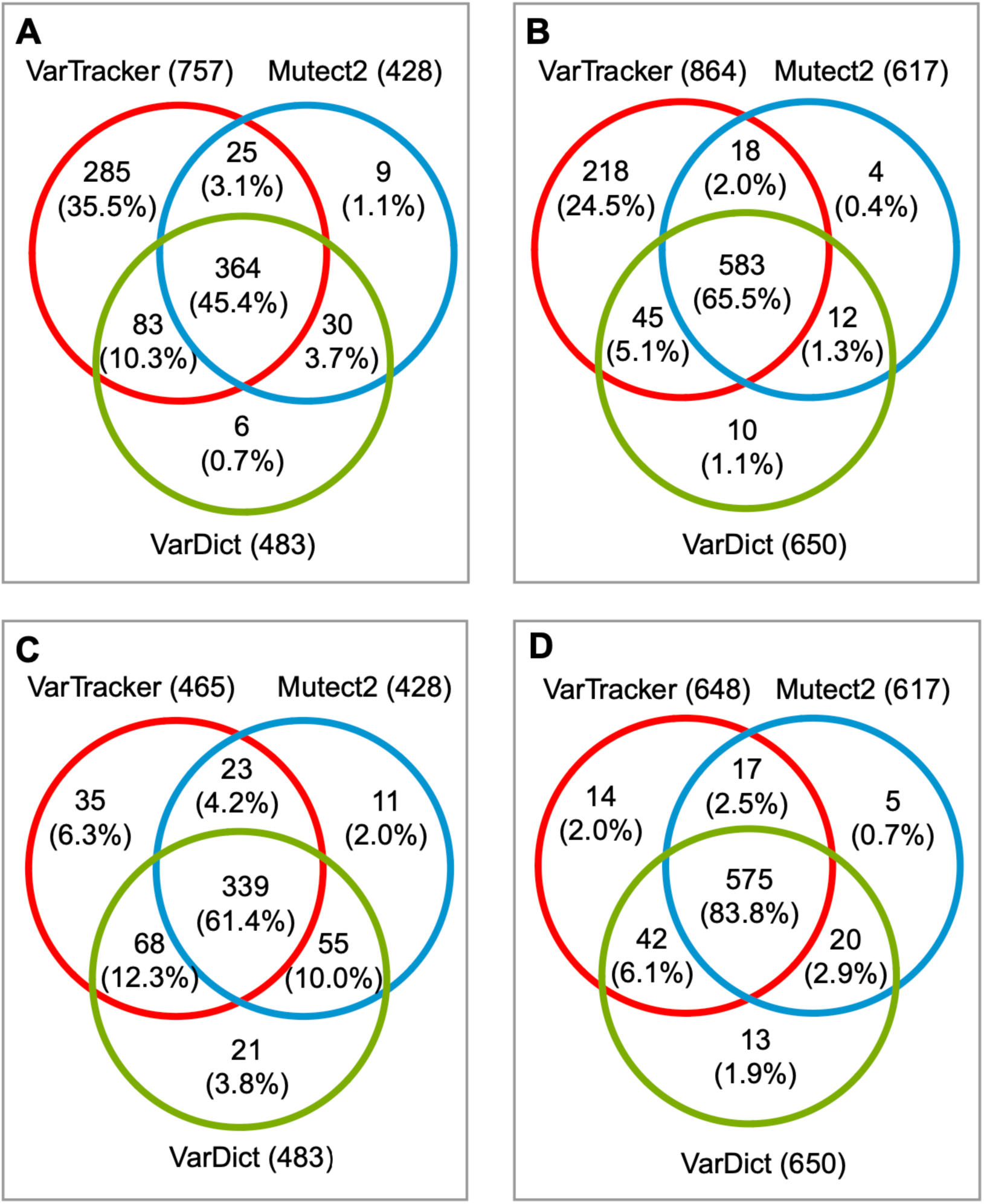
Overlap of spike-in CHIP recovered by the three callers. (**A**) NA12878_02 WES simulated at 100X coverage. At least one read supports alternative allele. (**B**) NA12878_02 WGS simulated at 50X coverage. At least one read supports alternative allele. (**C**) The same WES data as in (A). At least two reads support alternative allele. (**D**) The same WGS data as in (B). At least two reads support alternative allele. Both data were simulated with CHIP-specific VAFs. In each plot, SNVs and INDELs are combined. While VarTracker can report variants with a single supporting read, Mutect2 and VarDict only output those with ≥2 supporting reads.

### ML-based variant classification in simulated data

It remains challenging to distinguish CHIP mutations from germline variants solely based on VAFs, especially for those occurring in large, mutated clones. Furthermore, the three tools selected for meta-calling, VarTracker in particular, had high recall but low precision based on the simulated data (Supplementary Figure S4A and B). We hypothesized that, by learning the underlying features of known CHIP versus germline variants and artifacts, ML approaches could filter out most of the non-CHIP variants, thus minimizing the manual inspection needed in variant refinement. Toward this, we first assessed the predictive performance of 24 ML models on batch 2 data simulated with CHIP-specific VAFs (Supplementary Table S2), involving four ML algorithms, two variant types (SNVs and INDELs) and three data types (WGS, WES and combined data) (Supplementary Figure S1).

At default parameter setting, compared to the other two, XGBoost and Random forest showed higher accuracy, recall, precision and F1 scores across nearly all models in CHIP prediction (Table 1). For XGBoost and Random forest (and the other two as well), the predictive performance is clearly better on INDELs than on SNVs, showing 0.1-0.2 increase in F1 score, 17-33% increase in recall, and 6.2-14.4% increase in accuracy (Table 1). Not surprisingly, both methods showed better performance in WGS compared to WES, which is likely due to the greater bias observed in the latter (e.g., less uniform coverage stemmed from variation in capture efficiency and greater GC bias) (43). Consequently, it is known that WGS is more powerful in identifying exonic variants than WES (44). Nevertheless, hyperparameter tuning showed negligible gains over the default parameters in overall performance (Supplementary Figure S6A-C).

**Table 1.** Performance metrics of machine learning algorithms in predicting CHIP.

| Data | Algorithm | SNV |  |  |  | INDEL |  |  |  |
| --- | --- | --- | --- | --- | --- | --- | --- | --- | --- |
|  |  | F1 score | Recall (%) | Precision (%) | Accuracy (%) | F1 score | Recall (%) | Precision (%) | Accuracy (%) |
| WES | SVM | 0.642 | 50.6 | 87.7 | 75.2 | 0.898 | 86.2 | 93.8 | 90.1 |
|  | rpart | 0.65 | 50.2 | 92.2 | 75.1 | 0.927 | 50.2 | 92.9 | 92.7 |
|  | Random forest | 0.763 | 63 | 96.7 | 81.5 | 0.96 | 96.4 | 95.7 | 95.9 |
|  | XGBoost | 0.766 | 64.8 | 93.8 | 82.4 | 0.961 | 96.5 | 95.7 | 95.9 |
| WGS | SVM | 0.752 | 62.2 | 95.2 | 81 | 0.937 | 94.5 | 92.9 | 91.3 |
|  | rpart | 0.732 | 59.7 | 94.6 | 79.8 | 0.944 | 95.5 | 93.3 | 91.3 |
|  | Random forest | 0.863 | 78.5 | 95.9 | 89.2 | 0.97 | 97.6 | 96.4 | 95.7 |
|  | XGBoost | 0.875 | 80.7 | 95.6 | 90.2 | 0.975 | 97.8 | 97.3 | 96.4 |
| WES+WGS | SVM | 0.691 | 55.4 | 91.8 | 77.6 | 0.914 | 90.4 | 92.4 | 90.3 |
|  | rpart | 0.681 | 52.4 | 97.1 | 76.2 | 0.929 | 94.7 | 91.1 | 91.2 |
|  | Random forest | 0.817 | 71 | 96.3 | 85.5 | 0.967 | 96.9 | 96.5 | 96 |
|  | XGBoost | 0.832 | 74.2 | 94.7 | 87 | 0.966 | 96.9 | 96.4 | 96.2 |

Similarly, XGBoost and Random forest also had higher accuracy, recall and precision in predicting germline SNVs; while in predicting germline INDELs, SVM also had comparable performance (Supplementary Figure S7A-F). Thus, XGBoost and Random forest models were selected as the variant classifiers, with the former having slightly higher F1 score (due to the increased recall) in SNV detection from WGS.

We also assessed the performance of neural network model on WGS data from batch 2 simulation with CHIP-specific VAFs. Compared to XGBoost and Random forest, the neural network model showed 7.3-16.1% decrease in recall, as well as 5.3-9.1% decrease in accuracy and precision (Supplementary Table S6). Thus, neural network was not implemented for CHIP prediction (Figure 1A).

In XGBoost model, WGS was moderately correlated with WES in SNV (Pearson correlation (r)=0.82) and INDEL (r=0.64) classification based on all features (Supplementary Table S4). To understand which of the features contribute most to the overall CHIP prediction and how their contributions vary by sequencing platform and variant type, we analyzed feature importance in XGBoost classifier. For simplicity, we plot the top 13 features, each having ≥0.02 gain in at least one of the variant sets (Figure 4A-D). Overall, Meta calling status (i.e., number of callers identifying a variant), mapping quality score and VAF represent the most important features across all the four variant sets, together achieving a gain of 0.54-0.74. Individually, the magnitude of contribution from each feature appears to be sequencing-and variant type-dependent. For example, in predicting INDELs (Figure 4D), mapping quality alone had >0.5 gain in WES, versus <0.2 in the other 3 variant sets. This observation recapitulates the essence of accurate alignments in INDEL discovery, especially for WES that is known to enrich with low-quality INDEL calls compared to WGS (45). In addition, Mutect2 calling status had larger contribution to variant classification in WGS compared to WES, supporting that the local assembly strategy implemented in Mutect2 has greater power in WGS due to its more uniform and continuous read coverage. Third, COSMIC match had noticeable contributions to the detection of SNV (gain >0.05), but much less for INDEL (gain <0.01), likely owing to the overrepresentation (∼7-fold) of SNVs in the database. On the other hand, homopolymer run, low complexity region and short tandem repeat impacted the classification of INDELs, but less so for that of SNVs (Figure 4C and D). Indeed, these three elements are known to be associated with high false positive calls (39,45) or high mutation rates of INDEL (46). Taken together, our analyses demonstrated that XGBoost models can reliably predict CHIP over a wide range of VAFs, and that the prediction power is primarily attributed to a few associated features of high importance.

**Figure 4.**
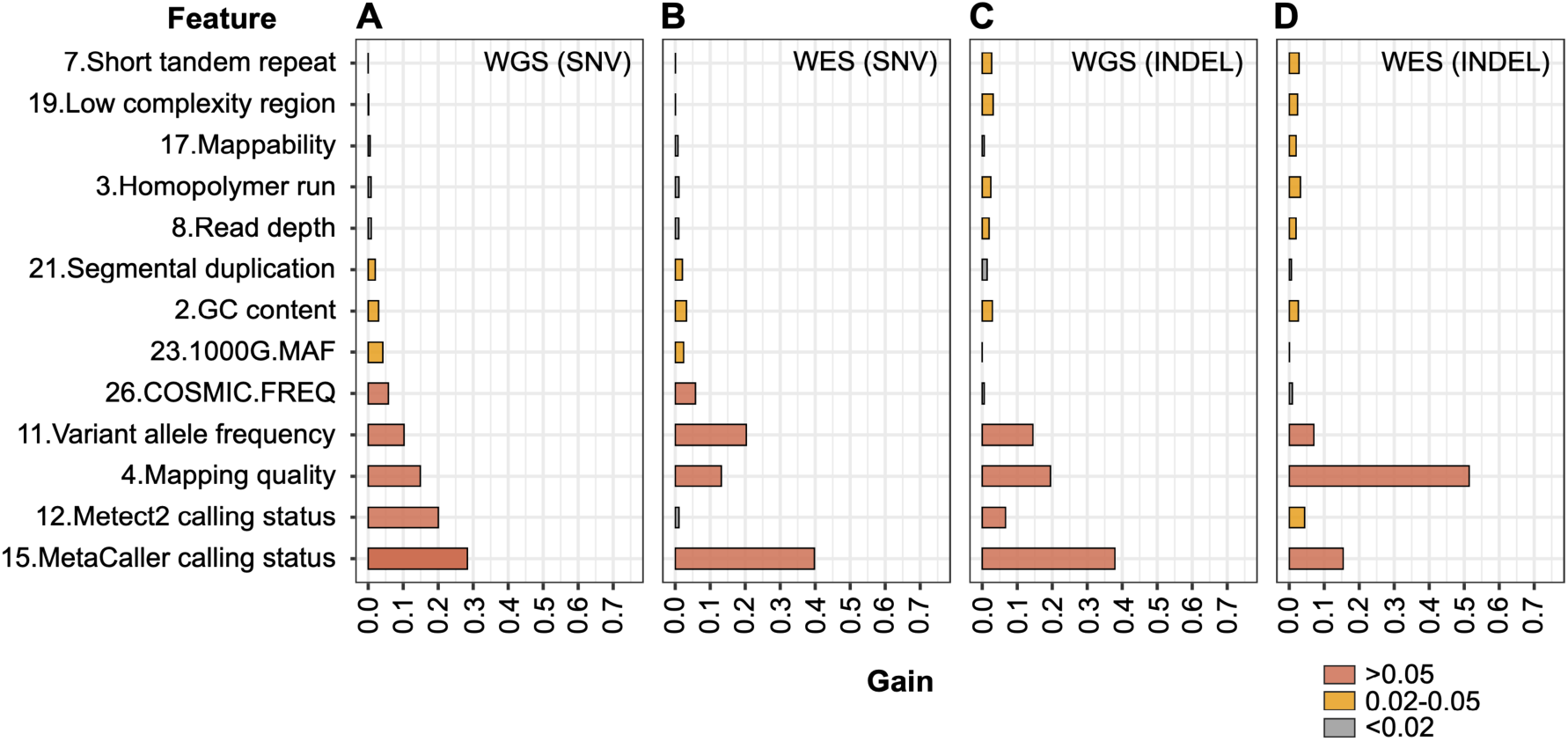
Contributions of the top 13 features to the predictive performance of XGBoost. (**A**) Feature’s gain in predicting SNV from WGS. (B) Prediction of SNV from WES. (**C**) Prediction of INDEL from WGS. (**D**) Prediction of INDEL from WES. The gain value is calculated using the xgb.importance function within the mIr package, which indicates a feature’s relative contribution to the model. The 13 features were selected to have gain values of ≥0.02 in at least one of the four predictions, showing in ascending order based on panel A. The number before each feature is from the “Feature” column in Supplementary Table S4.

### Pipeline validation with WES data

We applied UNISOM to real WES data from a cohort of 25 subjects confirmed to be absence of cytopenias and other hematologic disorders. To assess its performance, CHIP mutations were also identified from the same cohort with targeted deep sequencing (Supplementary Table S1). Specifically, from targeted panel, a total of 45 CHIP variants (only include SNVs due to very few INDELs) were retained after manual checking. In WES, 5 of these were mapped outside of the capture regions and another 7 had no coverage, leaving the other 33 (include 12 known CHIP) as ground truths.

WES data had an average of 158X coverage, with approximately 90% of the regions having at least 20X coverage. For each subject, meta-caller identified 18,080 raw variants on average, from which 11 were predicted as CHIP based on ML models trained on WES and WES+WGS, and 13 as CHIP based on WGS model. The vast majority (>99.9%) of the raw variants were classified as artifacts. The result indicates that, by automatically eliminating the artifacts, our approach could substantially reduce the manual work to refine CHIP candidates.

For the 33 CHIP mutations (“ground truths”) identified by targeted sequencing, the three models predicted 26-27 of them as CHIP in WES, including all the 12 known CHIP mutations previously reported (4,7,8), with 76.5-79.4% recall (Table 2). These CHIP mutations each had at least 2 supporting reads, with 5.6-51.4% VAF for 22 and 1.3-4.1% VAF for the other 5 mutations. The remaining six CHIP mutations were mis-predicted as artifacts; four the six were identified by VarTracker alone with a single read supporting, failing to meet the cutoff of ≥2 supporting reads, while the other two were identified by VarTracker and VarDict but had low (<3%) VAFs.

**Table 2.**
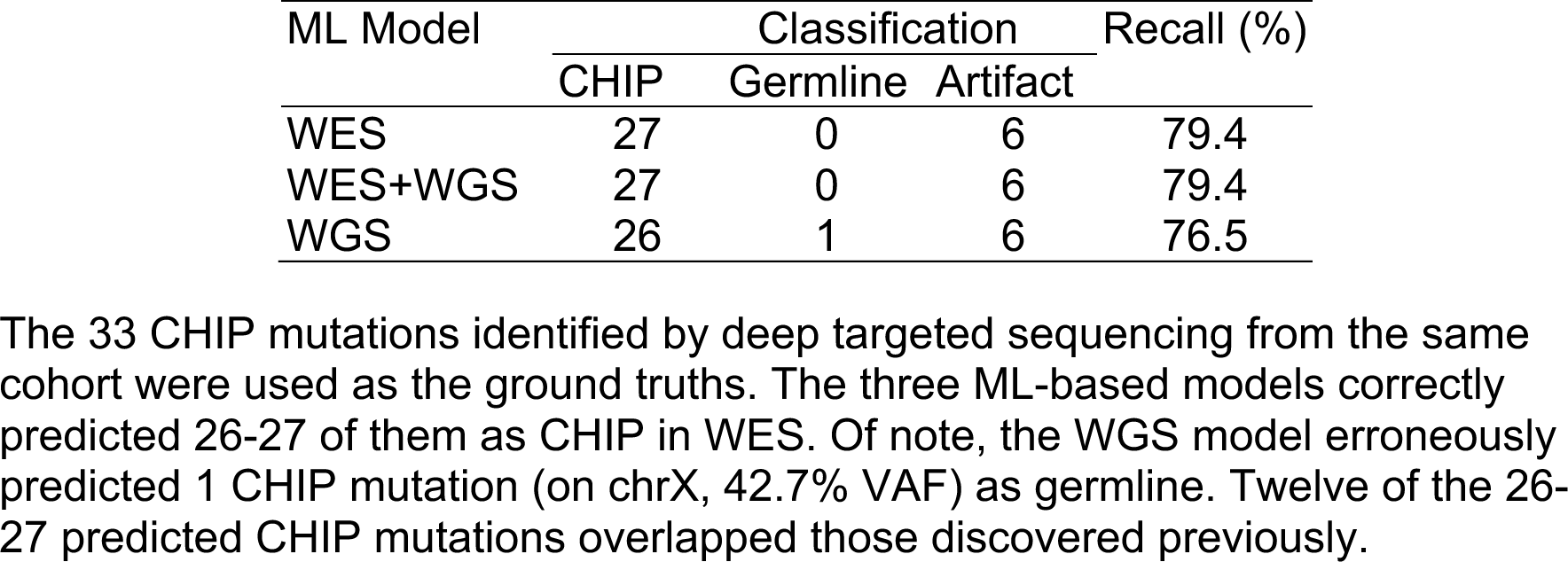
Prediction of CHIP in WES data.

| ML Model | Classification |  |  | Recall (%) |
| --- | --- | --- | --- | --- |
|  | CHIP | Germline | Artifact |  |
| WES | 27 | 0 | 6 | 79.4 |
| WES+WGS | 27 | 0 | 6 | 79.4 |
| WGS | 26 | 1 | 6 | 76.5 |

The 33 CHIP mutations identified by deep targeted sequencing from the same cohort were used as the ground truths. The three ML-based models correctly predicted 26-27 of them as CHIP in WES. Of note, the WGS model erroneously predicted 1 CHIP mutation (on chrX, 42.7% VAF) as germline. Twelve of the 26-27 predicted CHIP mutations overlapped those discovered previously.

### CHIP discovery in Mayo Clinic Biobank

To demonstrate UNISOM application in large-scale cohort studies, we analyzed WGS data (∼30X) from 979 participants in the Mayo Clinic Biobank who self-reported no diagnosis of hematological malignancies. Briefly, variants were identified using meta-caller at default parameter setting, followed by CHIP prediction on CAVA-annotated variants using the WGS-trained classifier. In comparison with previous CHIP studies using WES (7) and WGS (4), our analysis also focused on the leukemogenic driver mutations used in both studies (4,7).

A total of 202 CHIP mutations were identified in 35 driver genes at ≥2% VAF, with at least 2 reads supporting the alternative allele. All the 35 genes are included in the Mayo CHIP gene panel and TOPMed study (4). Two-thirds (133/202) represented mutations discovered in previous CHIP studies (4,7,8). We first examined the VAF distribution stratified by caller (Figure 5A). Both known and novel mutations had similar median VAF (5.6% versus 5.9%), which is about threefold lower than that from WGS-based TOPMed study (4). Looking closely at the clonal fraction, 84% of the 202 mutations had low VAFs of 2-10%. Among these low-VAF mutations, nearly 90% were identified by VarTracker alone or together with VarDict, but missed by Mutect2 (Figure 5A). The result supports the estimation that, for 30X WGS (equivalent to the average coverage in Mayo Biobank cohort), Mutect2 would miss about 90% of the CHIP mutations with 2-10% VAFs (4). Our analysis demonstrates the high sensitivity of the meta-calling strategy toward low VAF mutations even without high sequencing coverage.

**Figure 5.**
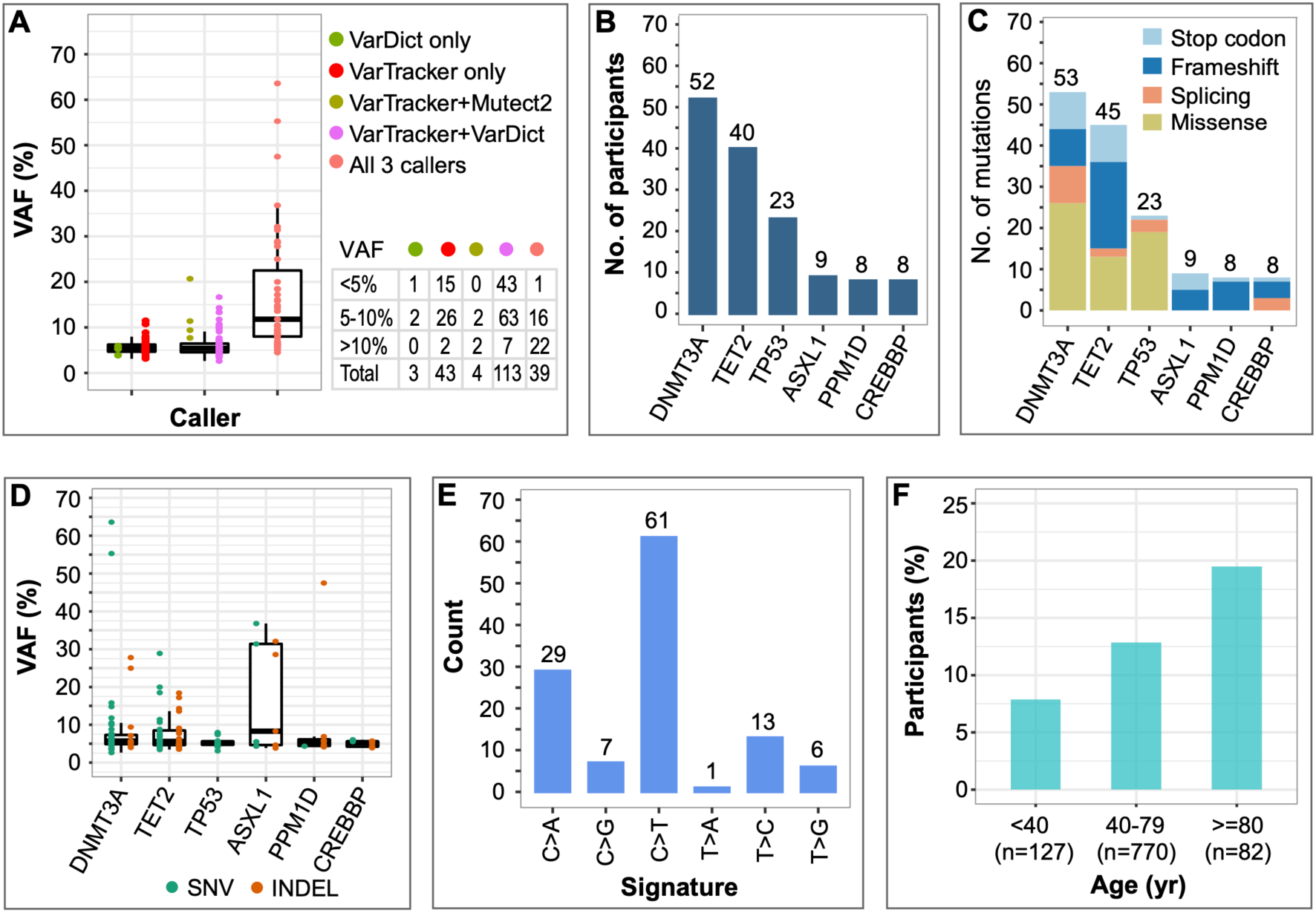
CHIP mutations in Mayo Biobank cohort. (**A**) VAFs of CHIP mutations. The bottom and upper lines in the box plot represent the 25% and 75% percentiles, with the horizontal line within representing the median. CHIP mutations were separated based on the tool(s) identifying them. No CHIP mutation was detected by Mutect2 alone, or only by Mutect2 and VarDict (“Mutect2+VarDict”). Of the three tools, VarTracker is most sensitive at low VAFs of ≤10%, with 41 unique calls. VarTracker and VarDict both identified CHIP with VAF down to 2.6%, while Mutect2 reached 4.5% VAF. (**B**) CHIP prevalence in the top six genes with the most mutations. For each gene, Y-axis shows the number of participants who carry at least one CHIP mutation in that gene. (**C**) Number of CHIP mutations in the six genes. The total mutations per gene were split by mutation type. For each gene, if two or more mutations occur in the same individual, or if the same mutation occurs in two or more individuals, they are summed up. (**D**) VAFs of CHIP mutations in the top six genes. No INDEL was identified in *TP53*. (**E**) Signatures enriched in CHIP mutations. (**F**) Prevalence of CHIP mutations in three age groups. There is a trend of increased CHIP prevalence by age as previously reported.

We next examined the CHIP prevalence in this cohort. Seventeen point five percent (171/979) of the participants carried CHIP mutations, which is much higher than the proportions reported in previous WES (7) and WGS studies (4). The high CHIP prevalence observed here is largely attributed to the meta-calling. Indeed, if only considering mutations identified by Mutect2 (Figure 5A), CHIP prevalence dropped to 3.9%, at a similar level (4.3%) as the TOPMed WGS project (∼40X) that used Mutect2 (4). Of the CHIP carriers, 143 individuals had a single mutation. The other 28 subjects each had 2-3 mutations in 16 genes (Supplementary Figure S8), ten of which previously were reported with co-mutations (7). Five of the 979 subjects did not provide the hematological phenotype status in the questionnaire. CHIP mutation was identified in only one of them, at 6% VAF within *EZH2*. Thus, the high CHIP prevalence observed in this cohort is unlikely due to the presence of cancer cells in blood.

CHIP mutations are known to be highly enriched in a few driver genes, including *DNMT3A*, *TET2*, and *ASXL1* that together harbored ∼70-80% of the mutations (4,7,8). Similar enrichment was observed in the Mayo Biobank cohort, with the top six genes carrying >70% of the mutations (Figure 5B and C). Five of them (except *CREBBP*) are among of the eight most frequently mutated genes (4,8). There was heterogeneity of clonal fraction within four of the six genes (*DNMT3A*, *TET2*, *ASXL1* and *PPM1D*) (Figure 5D). For example, the median clonal fraction in *ASXL1* (∼8.3%) is higher than that of the other five genes (5.2-5.7%), a trend previously reported (4).

We examined the type of mutations in the six genes (Figure 5C), which revealed similar portions of disruptive versus missense mutations, as previously reported (7,8). In *TP53*, majority (∼ 60-80%) of the mutations are missense mutations (7,47,48), a pattern we also revealed in this cohort. In contrast, these four genes (T*ET2*, *ASXL1*, *PPM1D* and *CREBBP*) predominantly or exclusively carried disruptive mutations, such as nonsense, frameshift and essential splice-site mutations (Figure 5C), consistent with previous findings (7,8). Finally, within *DNMT3A*, about half (51%) of the mutations are disruptive mutations, a proportion comparable to that (49%) from (7). As previously found in CHIP mutations (4,7), the most common single nucleotide substitution is cytosine to thymine (C>T) transition (52%), a signature known to be associated with aging (49,50), followed by cytosine to adenine (C>A) transversion and thymine to cytosine (T>C) transition (Figure 5E).

Lastly, we analyzed factors known to be associated with CHIP prevalence, including age (4,7,8) and smoking status (4,8). In this cohort, considering the six most commonly mutated genes, CHIP prevalence was strongly correlated with age at the time of sample collection (p-value=7.7e-03), ranging from 8% for subjects <40 years of age to nearly 20% among subjects ≥80 years of age (Figure 5F). Smoking was also associated with increased CHIP prevalence (odds ratio (OR) =1.47; 95% confidence interval 1.05-2.07; p-value=0.026) after adjusting for age in the multivariate logistic regression model. In summary, by applying the pipeline to the Mayo Biobank WGS data, our analyses revealed similar trends, as previously discovered in much large cohorts (4,7,8), in terms of most commonly mutated genes, type of mutations, mutation signature, as well as CHIP association with age and smoking status. In particular, the meta-calling approach markedly enhances the detection of low-VAF mutations, revealing high CHIP prevalence in this cohort with ∼30X coverage.

## DISCUSSION

In the general population without evidence of hematologic disorders, CHIP prevalence is estimated at ∼3-5% based on WES (7,8,51) and WGS data (4). However, limited by the coverage, sequencing error (>0.1%) (52) and the power of analysis pipeline, WGS and WES likely miss a high proportion of small mutant clones (5). To support this, smMIPs sequencing, which has an error rate of 10-fold lower compared to conventional sequencing, recovered over half of the CHIP missed by WGS data with ∼30X coverage (53). To tackle the analytical challenge, we present UNISOM as a standalone application for CHIP prediction. UNISOM utilizes a meta-calling strategy to achieve high sensitivity in variant detection, combined with a ML-based model for variant classification. Applied to the Mayo Biobank WGS data, we revealed a CHIP prevalence of 17.5% that is much higher compared to previous studies using a single caller (4,7), which is less sensitive to low-VAF mutations. Notably, over half of the clones carrying CHIP mutations continue to grow (16,53). For example, samples taken from individuals ∼6 years before diagnosed with AML were found to accumulate more mutations and show greater clonal expansion in peripheral blood compared to the control (16). Thus, implementing sensitive tools like UNISOM will improve the early detection of low-VAF CHIP mutations prior to diagnosis, enabling monitoring of the carriers and possible intervention (15,51).

Furthermore, analysis of longitudinal data began to shed insight into the dynamics of small clones (54). Variant fitness effect model predicted that over 80% of the variants with ≥1% VAF continue expansion over time (55). Consequently, some of the growing clones with ≥1% VAF, initially not considered as CHIP, will reach ≥2% VAF years later (53,55). It has been reported that clonal mutations with ≥1% VAF also confer an increased risk of developing AML (6). Although clinical implications of small clones remain unclear (5), the possibility of their expansion into larger clones suggests that close monitoring should be considered. Currently, UNISOM is limited to the detection of somatic mutations in leukemia driver genes. The applicability of UNISOM to other genes requires further testing.

UNISOM is sensitive even at insufficient sequencing coverage, as in the case of Mayo Biobank WGS data. UNISOM identified 30% of the CHIP mutations with 2-5% VAFs and another 54% with 5-10% VAFs, at a lenient cutoff of ≥2 supporting reads. The power of our approach is well illustrated by the CHIP mutations identified in *TP53* (Figure 5D). Not just in cancers (56), *TP53* mutations are also prevalent in blood cells of healthy individuals (57). Based on error-corrected targeted sequencing data, about half of the cancer-free elderly individuals carry rare *TP53* mutations with 0.01-0.37% VAFs (57). A similar phenomenon was also observed in patients with ovarian cancer and without cancer, who carried mutations with 0.01-0.1% VAFs that were predicted as being likely functional (58). Thus, low-VAF somatic mutations in *TP53* are unlikely to be noise. In the Mayo Biobank, *TP53* is the third most frequently mutated gene, with a total of 18 (12 known) CHIP mutations at 3.1-8.0% VAFs. All these mutations were present in exons 5-9, consistent with what has been reported that *TP53* mutations are predominantly clustered in exons 4-10 (58). Seven of the 12 known CHIP mutations overlap those identified in the TOPMed WGS data (∼40X) (4), but they were 2.3- to 5.1-fold lower in VAF. Not surprisingly, none of the 18 mutations were detected by Mutect2 that was found to be highly insensitive on 30X WGS data (4).

## DATA AVAILABILITY

All scripts for Meta-caller and ML classifiers are deposited in GitHub repository at https://github.com/shulanmayo/UNISOM.

## ACKNOWLEDGEMENTS

The analyses were performed in The National Center for Supercomputing Applications (https://www.ncsa.illinois.edu). This research was supported by the Mayo Clinic Center for Individualized Medicine.

## FUNDING

This work was supported by the Mayo Clinic Center for Individualized Medicine.

## Conflict of interest statement

None declared.

## Notes

### Competing Interest Statement

The authors have declared no competing interest.

## REFERENCES

1. Steensma, D.P., Bejar, R., Jaiswal, S., Lindsley, R.C., Sekeres, M.A., Hasserjian, R.P. and Ebert, B.L. (2015) Clonal hematopoiesis of indeterminate potential and its distinction from myelodysplastic syndromes. Blood, 126, 9–16.

2. Steensma, D.P. (2018) Clinical Implications of Clonal Hematopoiesis. Mayo Clin. Proc., 93, 1122–1130.

3. Khetarpal, S.A., Qamar, A., Bick, A.G., Fuster, J.J., Kathiresan, S., Jaiswal, S. and Natarajan, P. (2019) Clonal Hematopoiesis of Indeterminate Potential Reshapes Age-Related CVD JACC Review Topic of the Week. J. Am. Coll. Cardiol., 74, 578–586.

4. Bick, A.G., Weinstock, J.S., Nandakumar, S.K., Fulco, C.P., Bao, E.L., Zekavat, S.M., Szeto, M.D., Liao, X., Leventhal, M.J., Nasser, J. et al. (2020) Inherited causes of clonal haematopoiesis in 97,691 whole genomes. Nature, 586, 763–768.

5. Jaiswal, S. and Ebert, B.L. (2019) Clonal hematopoiesis in human aging and disease. Science, 366.

6. Young, A.L., Tong, R.S., Birmann, B.M. and Druley, T.E. (2019) Clonal hematopoiesis and risk of acute myeloid leukemia. Haematologica, 104, 2410–2417.

7. Jaiswal, S., Fontanillas, P., Flannick, J., Manning, A., Grauman, P.V., Mar, B.G., Lindsley, R.C., Mermel, C.H., Burtt, N., Chavez, A. et al. (2014) Age-related clonal hematopoiesis associated with adverse outcomes. N. Engl. J. Med., 371, 2488–2498.

8. Genovese, G., Kahler, A.K., Handsaker, R.E., Lindberg, J., Rose, S.A., Bakhoum, S.F., Chambert, K., Mick, E., Neale, B.M., Fromer, M. et al. (2014) Clonal hematopoiesis and blood-cancer risk inferred from blood DNA sequence. N. Engl. J. Med., 371, 2477–2487.

9. Jaiswal, S., Natarajan, P., Silver, A.J., Gibson, C.J., Bick, A.G., Shvartz, E., McConkey, M., Gupta, N., Gabriel, S., Ardissino, D. et al. (2017) Clonal Hematopoiesis and Risk of Atherosclerotic Cardiovascular Disease. N. Engl. J. Med., 377, 111–121.

10. Gibson, C.J., Lindsley, R.C., Tchekmedyian, V., Mar, B.G., Shi, J., Jaiswal, S., Bosworth, A., Francisco, L., He, J., Bansal, A. et al. (2017) Clonal Hematopoiesis Associated With Adverse Outcomes After Autologous Stem-Cell Transplantation for Lymphoma. J. Clin. Oncol., 35, 1598–1605.

11. Mouhieddine, T.H., Sperling, A.S., Redd, R., Park, J., Leventhal, M., Gibson, C.J., Manier, S., Nassar, A.H., Capelletti, M., Huynh, D. et al. (2020) Clonal hematopoiesis is associated with adverse outcomes in multiple myeloma patients undergoing transplant. Nat. Commun., 11, 2996.

12. Coombs, C.C., Zehir, A., Devlin, S.M., Kishtagari, A., Syed, A., Jonsson, P., Hyman, D.M., Solit, D.B., Robson, M.E., Baselga, J. et al. (2017) Therapy-Related Clonal Hematopoiesis in Patients with Non-hematologic Cancers Is Common and Associated with Adverse Clinical Outcomes. Cell Stem Cell, 21, 374–382.e374.

13. Acuna-Hidalgo, R., Sengul, H., Steehouwer, M., van de Vorst, M., Vermeulen, S.H., Kiemeney, L., Veltman, J.A., Gilissen, C. and Hoischen, A. (2017) Ultra-sensitive Sequencing Identifies High Prevalence of Clonal Hematopoiesis-Associated Mutations throughout Adult Life. Am. J. Hum. Genet., 101, 50–64.

14. Young, A.L., Challen, G.A., Birmann, B.M. and Druley, T.E. (2016) Clonal haematopoiesis harbouring AML-associated mutations is ubiquitous in healthy adults. Nat. Commun., 7.

15. Desai, P., Mencia-Trinchant, N., Savenkov, O., Simon, M.S., Cheang, G., Lee, S., Samuel, M., Ritchie, E.K., Guzman, M.L., Ballman, K.V. et al. (2018) Somatic mutations precede acute myeloid leukemia years before diagnosis. Nat. Med., 24, 1015–1023.

16. Abelson, S., Collord, G., Ng, S.W.K., Weissbrod, O., Mendelson Cohen, N., Niemeyer, E., Barda, N., Zuzarte, P.C., Heisler, L., Sundaravadanam, Y. et al. (2018) Prediction of acute myeloid leukaemia risk in healthy individuals. Nature, 559, 400–404.

17. Wang, Q., Jia, P., Li, F., Chen, H., Ji, H., Hucks, D., Dahlman, K.B., Pao, W. and Zhao, Z. (2013) Detecting somatic point mutations in cancer genome sequencing data: a comparison of mutation callers. Genome Med., 5, 91.

18. Xu, H., DiCarlo, J., Satya, R.V., Peng, Q. and Wang, Y. (2014) Comparison of somatic mutation calling methods in amplicon and whole exome sequence data. BMC Genomics, 15, 244.

19. Krøigård, A.B., Thomassen, M., Lænkholm, A.V., Kruse, T.A. and Larsen, M.J. (2016) Evaluation of Nine Somatic Variant Callers for Detection of Somatic Mutations in Exome and Targeted Deep Sequencing Data. PLoS One, 11, e0151664.

20. Cai, L., Yuan, W., Zhang, Z., He, L. and Chou, K.C. (2016) In-depth comparison of somatic point mutation callers based on different tumor next-generation sequencing depth data. Sci. Rep., 6, 36540.

21. Fang, L.T., Afshar, P.T., Chhibber, A., Mohiyuddin, M., Fan, Y., Mu, J.C., Gibeling, G., Barr, S., Asadi, N.B., Gerstein, M.B. et al. (2015) An ensemble approach to accurately detect somatic mutations using SomaticSeq. Genome Biol., 16, 197.

22. Bailey, M.H., Meyerson, W.U., Dursi, L.J., Wang, L.B., Dong, G., Liang, W.W., Weerasinghe, A., Li, S., Li, Y., Kelso, S. et al. (2020) Retrospective evaluation of whole exome and genome mutation calls in 746 cancer samples. Nat. Commun., 11, 4748.

23. Ellrott, K., Bailey, M.H., Saksena, G., Covington, K.R., Kandoth, C., Stewart, C., Hess, J., Ma, S., Chiotti, K.E., McLellan, M. et al. (2018) Scalable Open Science Approach for Mutation Calling of Tumor Exomes Using Multiple Genomic Pipelines. Cell Syst., 6, 271–281.e277.

24. Campbell, P.J., Getz, G., Korbel, J.O., Stuart, J.M., Jennings, J.L., Stein, L.D., Perry, M.D., Nahal-Bose, H.K., Ouellette, B.F.F., Li, C.H. et al. (2020) Pan-cancer analysis of whole genomes. Nature, 578, 82–93.

25. Dou, Y., Gold, H.D., Luquette, L.J. and Park, P.J. (2018) Detecting Somatic Mutations in Normal Cells. Trends Genet., 34, 545–557.

26. Xie, M., Lu, C., Wang, J., McLellan, M.D., Johnson, K.J., Wendl, M.C., McMichael, J.F., Schmidt, H.K., Yellapantula, V., Miller, C.A. et al. (2014) Age-related mutations associated with clonal hematopoietic expansion and malignancies. Nat. Med., 20, 1472–1478.

27. Niroula, A., Sekar, A., Murakami, M.A., Trinder, M., Agrawal, M., Wong, W.J., Bick, A.G., Uddin, M.M., Gibson, C.J., Griffin, G.K. et al. (2021) Distinction of lymphoid and myeloid clonal hematopoiesis. Nat. Med., 27, 1921–1927.

28. Zook, J.M., Chapman, B., Wang, J., Mittelman, D., Hofmann, O., Hide, W. and Salit, M. (2014) Integrating human sequence data sets provides a resource of benchmark SNP and indel genotype calls. Nat. Biotechnol., 32, 246–251.

29. Zook, J.M., McDaniel, J., Olson, N.D., Wagner, J., Parikh, H., Heaton, H., Irvine, S.A., Trigg, L., Truty, R., McLean, C.Y. et al. (2019) An open resource for accurately benchmarking small variant and reference calls. Nat. Biotechnol., 37, 561–566.

30. Ewing, A.D., Houlahan, K.E., Hu, Y., Ellrott, K., Caloian, C., Yamaguchi, T.N., Bare, J.C., P’ng, C., Waggott, D., Sabelnykova, V.Y. et al. (2015) Combining tumor genome simulation with crowdsourcing to benchmark somatic single-nucleotide-variant detection. Nat. Methods, 12, 623–630.

31. Li, H., Handsaker, B., Wysoker, A., Fennell, T., Ruan, J., Homer, N., Marth, G., Abecasis, G. and Durbin, R. (2009) The Sequence Alignment/Map format and SAMtools. Bioinformatics, 25, 2078–2079.

32. Stachler, M.D., Taylor-Weiner, A., Peng, S., McKenna, A., Agoston, A.T., Odze, R.D., Davison, J.M., Nason, K.S., Loda, M., Leshchiner, I. et al. (2015) Paired exome analysis of Barrett’s esophagus and adenocarcinoma. Nat. Genet., 47, 1047–1055.

33. Xu, C. (2018) A review of somatic single nucleotide variant calling algorithms for next-generation sequencing data. Comput. Struct. Biotechnol. J., 16, 15–24.

34. Münz, M., Ruark, E., Renwick, A., Ramsay, E., Clarke, M., Mahamdallie, S., Cloke, V., Seal, S., Strydom, A., Lunter, G. et al. (2015) CSN and CAVA: variant annotation tools for rapid, robust next-generation sequencing analysis in the clinical setting. Genome Med., 7, 76.

35. Bischl, B., Lang, M., Kotthoff, L., Schiffner, J., Richter, J., Studerus, E., Casalicchio, G. and Jones, Z.M. (2016) mlr: Machine Learning in R. J. Mach. Learn. Res., 17, 1–5.

36. Kraft, I.L. and Godley, L.A. (2020) Identifying potential germline variants from sequencing hematopoietic malignancies. Blood, 136, 2498–2506.

37. Olson, J.E., Ryu, E., Johnson, K.J., Koenig, B.A., Maschke, K.J., Morrisette, J.A., Liebow, M., Takahashi, P.Y., Fredericksen, Z.S., Sharma, R.G. et al. (2013) The Mayo Clinic Biobank: a building block for individualized medicine. Mayo Clin. Proc., 88, 952–962.

38. Pottier, C., Ren, Y., Perkerson, R.B., 3rd, Baker, M., Jenkins, G.D., van Blitterswijk, M., DeJesus-Hernandez, M., van Rooij, J.G.J., Murray, M.E., Christopher, E. et al. (2019) Genome-wide analyses as part of the international FTLD-TDP whole-genome sequencing consortium reveals novel disease risk factors and increases support for immune dysfunction in FTLD. Acta Neuropathol., 137, 879–899.

39. Li, H. (2014) Toward better understanding of artifacts in variant calling from high-coverage samples. Bioinformatics, 30, 2843–2851.

40. Dorsheimer, L., Assmus, B., Rasper, T., Ortmann, C.A., Ecke, A., Abou-El-Ardat, K., Schmid, T., Brüne, B., Wagner, S., Serve, H. et al. (2019) Association of Mutations Contributing to Clonal Hematopoiesis With Prognosis in Chronic Ischemic Heart Failure. JAMA Cardiol., 4, 25–33.

41. Said Mohammed, K., Kibinge, N., Prins, P., Agoti, C.N., Cotten, M., Nokes, D.J., Brand, S. and Githinji, G. (2018) Evaluating the performance of tools used to call minority variants from whole genome short-read data. Wellcome Open Res., 3, 21.

42. Tian, S., Yan, H., Neuhauser, C. and Slager, S.L. (2016) An analytical workflow for accurate variant discovery in highly divergent regions. BMC Genomics, 17, 703.

43. Meienberg, J., Bruggmann, R., Oexle, K. and Matyas, G. (2016) Clinical sequencing: is WGS the better WES? Hum. Genet., 135, 359–362.

44. Belkadi, A., Bolze, A., Itan, Y., Cobat, A., Vincent, Q.B., Antipenko, A., Shang, L., Boisson, B., Casanova, J.L. and Abel, L. (2015) Whole-genome sequencing is more powerful than whole-exome sequencing for detecting exome variants. Proc. Natl. Acad. Sci. U. S. A., 112, 5473–5478.

45. Fang, H., Wu, Y., Narzisi, G., O’Rawe, J.A., Barrón, L.T., Rosenbaum, J., Ronemus, M., Iossifov, I., Schatz, M.C. and Lyon, G.J. (2014) Reducing INDEL calling errors in whole genome and exome sequencing data. Genome Med., 6, 89.

46. Maruvka, Y.E., Mouw, K.W., Karlic, R., Parasuraman, P., Kamburov, A., Polak, P., Haradhvala, N.J., Hess, J.M., Rheinbay, E., Brody, Y. et al. (2017) Analysis of somatic microsatellite indels identifies driver events in human tumors. Nat. Biotechnol., 35, 951–959.

47. Shirole, N.H., Pal, D., Kastenhuber, E.R., Senturk, S., Boroda, J., Pisterzi, P., Miller, M., Munoz, G., Anderluh, M., Ladanyi, M. et al. (2016) TP53 exon-6 truncating mutations produce separation of function isoforms with pro-tumorigenic functions. Elife, 5.

48. Boettcher, S., Miller, P.G., Sharma, R., McConkey, M., Leventhal, M., Krivtsov, A.V., Giacomelli, A.O., Wong, W., Kim, J., Chao, S. et al. (2019) A dominant-negative effect drives selection of TP53 missense mutations in myeloid malignancies. Science, 365, 599–604.

49. Welch, J.S., Ley, T.J., Link, D.C., Miller, C.A., Larson, D.E., Koboldt, D.C., Wartman, L.D., Lamprecht, T.L., Liu, F., Xia, J. et al. (2012) The origin and evolution of mutations in acute myeloid leukemia. Cell, 150, 264–278.

50. Alexandrov, L.B., Nik-Zainal, S., Wedge, D.C., Aparicio, S.A.J.R., Behjati, S., Biankin, A.V., Bignell, G.R., Bolli, N., Borg, A., Børresen-Dale, A.-L. et al. (2013) Signatures of mutational processes in human cancer. Nature, 500, 415–421.

51. Feusier, J.E., Arunachalam, S., Tashi, T., Baker, M.J., VanSant-Webb, C., Ferdig, A., Welm, B.E., Rodriguez-Flores, J.L., Ours, C., Jorde, L.B. et al. (2021) Large-Scale Identification of Clonal Hematopoiesis and Mutations Recurrent in Blood Cancers. Blood Cancer Discov., 2, 226–237.

52. Ma, X., Shao, Y., Tian, L., Flasch, D.A., Mulder, H.L., Edmonson, M.N., Liu, Y., Chen, X., Newman, S., Nakitandwe, J. et al. (2019) Analysis of error profiles in deep next-generation sequencing data. Genome Biol., 20, 50.

53. Uddin, M.M., Zhou, Y., Bick, A.G., Burugula, B.B., Jaiswal, S., Desai, P., Honigberg, M.C., Love, S.A., Barac, A., Hayden, K.M. et al. (2022) Longitudinal profiling of clonal hematopoiesis provides insight into clonal dynamics. Immun. Ageing, 19, 23.

54. Evans, M.A. and Walsh, K. (2023) Clonal hematopoiesis, somatic mosaicism, and age-associated disease. Physiol. Rev., 103, 649–716.

55. Robertson, N.A., Latorre-Crespo, E., Terradas-Terradas, M., Lemos-Portela, J., Purcell, A.C., Livesey, B.J., Hillary, R.F., Murphy, L., Fawkes, A., MacGillivray, L. et al. (2022) Longitudinal dynamics of clonal hematopoiesis identifies gene-specific fitness effects. Nat. Med., 28, 1439–1446.

56. Hu, J., Cao, J., Topatana, W., Juengpanich, S., Li, S., Zhang, B., Shen, J., Cai, L., Cai, X. and Chen, M. (2021) Targeting mutant p53 for cancer therapy: direct and indirect strategies. J. Hematol. Oncol., 14, 157.

57. Wong, T.N., Ramsingh, G., Young, A.L., Miller, C.A., Touma, W., Welch, J.S., Lamprecht, T.L., Shen, D., Hundal, J., Fulton, R.S. et al. (2015) Role of TP53 mutations in the origin and evolution of therapy-related acute myeloid leukaemia. Nature, 518, 552–555.

58. Krimmel, J.D., Schmitt, M.W., Harrell, M.I., Agnew, K.J., Kennedy, S.R., Emond, M.J., Loeb, L.A., Swisher, E.M. and Risques, R.A. (2016) Ultra-deep sequencing detects ovarian cancer cells in peritoneal fluid and reveals somatic TP53 mutations in noncancerous tissues. Proc. Natl. Acad. Sci. U. S. A., 113, 6005–6010.

