## Supplement1 for "Unified somatic calling and machine learning-based classification enhance the discovery of clonal hematopoiesis of indeterminate potential"

**Supplementary note 1**

### Unified somatic calling and machine learning-based classification enhance the discovery of clonal hematopoiesis of indeterminate potential

Shulan Tian, Garrett Jenkinson, Alejandro Ferrer, Huihuang Yan, Joel A. Morales-Rosado, Kevin L. Wang, Terra L. Lasho, Benjamin B. Yan, Saurabh Baheti, Janet E. Olson, Linda B. Baughn, Wei Ding, Susan L. Slager, Mrinal S. Patnaik, Konstantinos N. Lazaridis, and Eric W. Klee

#### Known CHIP mutations

A list of 2,367 CHIP mutations, including 1,331 SNVs and 1,036 INDELs (Supplementary Table S1), was compiled from two WES-based CHIP studies (1,2) and the WGS-based TOPMed study (3). The analysis was based on the following rules: (1) SNVs were prioritized over INDELs if both were reported at the same genomic position across these studies or across different subjects in the same study; (2) for variants identified at the same genomic position in multiple subjects, the one with the lowest VAF was retained along with its alternate base in that subject; (3) for a given variant, if two or three alternate bases were identified at the same VAF, one of them was randomly selected; and (4) INDELs were normalized as left-aligned and parsimonious with GATK walker LeftAlignAndTrimVariants. In the resulting list, over 92% of the CHIP mutations were identified from WGS in the TOPMed cohort that has the largest sample size with nearly 97,700 participants (3).

#### ML-based variant classifier

CHIP mutations often had low VAFs, in the same range as artifacts from DNA contamination and damage, or from sequencing and alignment errors, which makes VAF-based filtering problematic. Also, due to the much lower somatic mutation rates in normal cells, artifacts are often observed much more frequently than true somatic variants. Thus, the filtering process is very critical to refine true CHIP mutations but is poorly standardized and time-consuming.

The traditional hard-filtering approach is based on attributes from variant annotations, such as sequence context and the number of reads supporting alternative allele. It performs a multiple-step filtering based on the pre-defined thresholds of individual attributes. With the challenges of setting optimal thresholds across all selected attributes simultaneously, this approach will filter out true mutations if they failed to meet any of the thresholds. In addition, CHIP mutations have characteristic mutational signatures (1,3), which probabilistically contribute to CHIP prediction but are difficult to be incorporated into the hard-filtering approach. On the other hand, ML-based models learn information from annotation profiles in the training set that can be used to discriminate real somatic mutations (CHIP) from germline variants and artifacts. Based on the assessment made on the test set, the best-performing ML model(s) will be selected and further applied to the real data for CHIP prediction. Unlike hard-filtering that requires multiple rounds of selections, ML-based probabilistic classifiers enable one-time binary selection of the predicted CHIP mutations. Thus, to streamline the CHIP detection, we developed ML-based prediction models based on variant-associated features.

To build ML-based prediction models, raw variants were pre-assigned with class labels, as described below, split into SNVs and INDELs, and the associated features were extracted. Four algorithms were each tested on SNVs and INDELs called from WES, WGS and the combined data (i.e., six tests per algorithm), at default parameter setting. In addition, neural network was tested on WGS data. In calculating the performance metrics, the pre-assigned class labels were used to represent the actual class labels. Based on the performance metrics, XGBoost and Random forest were selected for further hyperparameter tuning. Key steps are illustrated in Supplementary Figure S1 and described below in detail.

***Class labels***

Variants from meta-calling were each assigned with an actual class label of being CHIP, GERMLINE, or ARTIFACT, To identify those that are most likely germline variants, we used a set of SNVs and short INDELs with high-confidence in NA12878 (NISTv3.3.2) as the ground truths (4), which is generated by the GIAB Consortium and publicly available at https://ftp-trace.ncbi.nlm.nih.gov/ReferenceSamples/giab/release/NA12878_HG001/latest/GRCh37/SupplementaryFiles/HG001_GRCh37_1_22_v4.2.1_all.vcf.gz. A variant is flagged as GERMLINE if (1) it overlaps the above GIAB list of known germline variants; or (2) it is a novel one but identified by at least two of the callers, with >30% VAF, a cutoff of being germline as previously suggested (2,5). The second criterion for including novel ons considers a possibility that the GIAB list may still miss some germline variants. On the other hand, a variant is flagged as CHIP if (1) it is mapped to the genomic position carrying a CHIP spike-in; or (2) it is called from elsewhere by at least two of the callers with ≤30% VAF, considering that NA12878 itself may also carry CHIP. The remaining variants are labeled as ARTIFACT. They were used as the actual class labels in calculating performance metrics on the test set for CHIP prediction models.

***Feature extraction***

To build a well-performing variant classifier, it is critical to select the features that together can potentially characterize a raw variant, so that the algorithm can learn the underlying discriminatory patterns differentiating between CHIP, GERMLINE, and ARTIFACT. For each raw SNV and INDEL with CAVA predicted functional effects, we collected 25 and 23 features, respectively, that describe meta-calling status, variant quality metrics, the associated genomic context, as well as the status in five public variant databases (Supplementary Table S4**)**. Additionally, we included mutational signature, considering that, in CH, >50% of the observed point mutations are C-to-T transitions (1,2), which are the dominant point mutations in most aging tissues (1,6). GATK walkers VariantAnnotator and VariantFiltration were used to extract these features.

***Model training***

Given the raw variants in the feature space, variant classifier predicts which ones are real CHIP with functional effects. As calling accuracy of INDEL remains lower than that of SNV (7), the two were separated in model training. Also, considering the possible differences in sequencing platforms, variant classifier was built separately for WES, WGS and the combined data. To avoid overfitting, for each of the six callsets (2 variant types x 3 data types), we applied the holdout method by randomly selecting 80% of the raw variants as the training set and leaving the remaining 20% as the test set.

To start model training, all 11 categorical variables were one-hot encoded into (Supplementary Table S4**)**. For each of the six callsets, the mlr R package (8) was used to build four ML models, including Recursive Partitioning and Regression Trees (rpart), Random forest, Support Vector Machine (SVM) and eXtreme Gradient Boosting

(XGBoost). We ran the four models on the training set, with the default parameters (Supplementary Table S5), and evaluated their performance in class prediction.

We also tested neural network in CHIP prediction, using WGS data from batch 2 simulation with CHIP-specific VAFs. It took the same 26 (SNV) and 24 (INDEL) features used to train the 4 ML-based models described above. Neural network model was built with TensorFlow platform (https://www.tensorflow.org) in python environment (Supplementary Table S5).

***Hyperparameter tuning***

XGBoost and Random forest based classifiers clearly demonstrated better performance in predicting CHIP compared to the other models. Thus, both were selected for hyperparameter tuning on the training set, aiming to optimize the model performance by adjusting various hyperparameters. Hyperparameter tuning used k-fold cross-validation (here k=5) (Supplementary Table S5), which was compared to the model built with default parameters.

**Confidence interval estimation**

To facilitate the prioritization, following the above prediction refinement, a hierarchical Bayesian model was developed to estimate confidence intervals for CHIP predictions. The model utilizes CHIP incidence in a reference population of healthy individuals as the background. It is known that CHIP mutations are very rare in young individuals under the age 40 (1,2). In numerous studies, healthy individuals in this age range were selected as normal controls when calling CHIP mutations from WGS (3) and WES data (9). To build the reference, WES data (median coverage 125X) were generated from 75 healthy individuals younger than 40 years of age. Under the model, CHIP is defined as the allele deviated from the reference one in this healthy population. Then, in an individual, the probability of detecting CHIP at a given position can be estimated based on the background model (Supplementary note 2).

**CHIP prediction with WES data**

The performance of meta-caller and XGBoost classifier was further evaluated on real WES data from a cohort of 25 individuals, for whom targeted deep sequencing data were also available. The study was approved by the institutional review board at Mayo Clinic

(Mayo Clinic IRB 16-004173). Peripheral blood mononuclear cells were collected, with written consent, from individuals who were examined by a medical professional for the absence of cytopenias, multiple myeloma and other hematologic disorders in blood. Bone marrow was also examined to verify the absence of multiple myeloma and lymphoma. DNA isolation, WES library construction, Illumina paired-end sequencing, reads mapping and post-alignment processing were performed as previously described (10). WES had a median coverage of 158X (127-179X), estimated by the GATK walker DepthOfCoverage. Variants were detected with meta-caller, with default parameter setting, followed by functional annotation with CAVA. Those predicted to have functional effects were used as input for the XGBoost classifier pretrained on simulated WES data.

To assess the CHIP predictions made by the XGBoost classifier, we used CHIP variants identified from targeted sequencing data as ground truths. In brief, DNA from the same 25 individuals with WES data was deeply sequenced on a customized panel of 189 leukemia-associated genes (Supplementary Table S1). Library construction, paired-end sequencing and bioinformatics analysis were as described (11). The 25 samples have approximately 95% of the targeted regions sequenced at ≥100X coverage, with ~50% up to 3,000-4,000X coverage and a median coverage of approximately 1,000X. At the aforementioned sequencing coverage, variants with >0.5% VAF should be identified with high confidence. Candidate mutations were identified by two somatic calling pipelines, an in-house pipeline (BWA-mem + GATK HaplotypeCaller) and the commercial Agilent SureCall (v4.1.2, BWA + SNPPET SNV caller), following the criteria used in (11). Variants were filtered out if present in any of the four public germline databases (the 1000 Genomes Project, gnomAD, ExAC and TOPMed; Supplementary Table S4) with ≥0.3% minor allele frequency (MAF).

The retained variants were classified as CHIP if they met any of the following criteria: (1) overlap with CHIP-associated driver mutations defined in (12); (2) present in COSMIC database; (3) predicted as being deleterious by SIFT (13), MutationTaster2 (14), or Polyphen-2 (15); or (4) with functional relevance in leukemia based on literature search. Finally, the identified CHIP mutations were manually inspected with Alamut Visual software (v2.7, Interactive Biosoftware, Rouen, Haute-Normandie, France), by which potential sequencing artifacts and those not in the list of known driver mutations were filtered out.

After the above manual curation, 45 CHIP mutations were retained. Those mapped outside of the WES capture regions (5 mutations) or with no coverage in WES (7 mutations) were filtered out. The remaining 33 mutations (45-5-7) were used as ground truths when assessing XGBoost predictions on WES from the same cohort. We used recall, based on formula (2), to measure the predictive performance, where true positive and false negative are those from the 33 mutations that are detected and missed by UNISOM, respectively.

**Supplementary Table S1**. Collection of leukemia-associated genes and known CHIP mutations

| Data | Type | No. gene | No. SNV | No. INDEL | No. subject | Ref. |
|---|---|---|---|---|---|---|
| 1 | WES | 156 | 310 | 181 | 17,182 | (1) |
| 2 | WES | 14 | 144 | 68 | 12,380 | (2) |
| 3* | WES | 31 | 65 | 12 | 2,728 | (16) |
| 4 | WGS | 74 | 1,168 | 1,082 | 97,691 | (3) |
| 5 | NA | 189 | NA | NA | NA | Mayo Clinic CHIP panel |
| Total | | 202 | 1,331 | 1,036 | | |

Genomic positions are not available for the 77 CHIP mutations (65 SNVs and 12 INDELs) identified in Xie_2014 (16). Mayo Clinic CHIP panel only has the gene list without variant information. Gene names with aliases from different studies are consolidated into a single one.

**Supplementary Table S2**. NA12878 WGS and WES data used in this study

| Data set | Type | Platform | Sample ID | Read length (bp) | Original coverage (X) | Coverage used (X) | Source* |
|---|---|---|---|---|---|---|---|
| 1 | WGS | HiSeq 2500 | GIAB_HG001_WGS | 148 | 300 | 200, 100, 50, 20 | GIAB (4) |
| 2 | WGS | HiSeq 4000 | SRR8454587 | 150 | 29 | 29 | SRA (17) |
| 3 | WGS | HiSeq 4000 | SRR8454588 | 150 | 29 | 29 | SRA (17) |
| 4 | WGS | NovaSeq 6000 | SRR8454589 | 150 | 29 | 29 | SRA (17) |
| 5** | WGS | HiSeq X Ten | SRR6885087 | 150 | 37 | 37 | SRA |
| 6** | WGS | HiSeq X Ten | SRR7733437 | 150 | 24 | 24 | SRA |
| 7 | WGS | HiSeq X Ten | SRR7781427 | 150 | 34 | 34 | SRA |
| 8 | WGS | HiSeq X Ten | SRR7781429 | 150 | 40 | 40 | SRA |
| 9 | WGS | HiSeq X Ten | SRR7781444 | 150 | 37 | 37 | SRA (17) |
| 10 | WGS | HiSeq X Ten | SRR7781431 | 150 | 24 | 24 | SRA (17) |
| 11 | WGS | NovaSeq | NA12878_01_WGS | 150 | 100 | 100, 50, 20 | Internal |
| 12 | WGS | NovaSeq | NA12878_02_WGS | 150 | 84 | 84, 50, 20 | Internal |
| 13 | WES | HiSeq 2500 | GIAB_HG001_WES | 100 | 100 | 100 | GIAB (4) |
| 14 | WES | HiSeq 4000 | SRR8381428 | 148 | 355 | 200, 100, 50, 20 | SRA |
| 15 | WES | NovaSeq 6000 | SRR8381429 | 145 | 213 | 200, 100, 50, 20 | SRA (17) |
| 16 | WES | HiSeq 4000 | ERR1905889 | 150 | 318 | 200, 100, 50, 20 | SRA |
| 17 | WES | HiSeq 4000 | ERR1905890 | 150 | 360 | 200, 100, 50, 20 | SRA |
| 18 | WES | HiSeq 2500 | NA12878_01_WES | 102 | 132 | 132, 100, 50, 20 | Internal |
| 19 | WES | HiSeq 2500 | NA12878_02_WES | 102 | 135 | 135, 100, 50, 20 | Internal |

*Source: GIAB, Genome in a Bottle Consortium; SRA, Sequence Read Archive; Internal, Mayo Clinic. Sequence at SRA can be downloaded through the web link: https://trace.ncbi.nlm.nih.gov/Traces/sra/?run=accession, with accession from the "Sample ID" column. **Data 5 and 6 were generated from PCR-free libraries. Batch 1 simulation used 21 BAMs from five WGS (1, 3, 4, 11, and 12) and three WES data (13, 18 and 19), with 13 uniform and CHIP-specific VAFs. Batch 2 simulation used all 44 BAMs from the 19 datasets, with CHIP-specific VAFs.

**Supplementary Table S3**. Eleven tools benchmarked for single-sample variant detection

| Caller | Algorithm | Variant type | Version | Parameter setting | Ref. |
|---|---|---|---|---|---|
| LoFreq | Sequencing error modeling, Poisson-binomial | SNVs and INDELs | v2.1.4 | lofreq call-parallel --pp-threads n_threads -f ref.fa -o output.tmp.vcf input.bam;<br>bgzip output.tmp.vcf;<br>tabix -p vcf output.tmp.vcf.gz; tabix -h -f output.tmp.vcf.gz target.bed > output.vcf | (18) |
| VarDict | Supervised and unsupervised local realignments, a heuristic method | SNVs, MNPs, INDELs and SV | v2019.06.04-0 | vardict -G ref.fa -f 0.01 -N output -b input.bam -c 1 -S 2 -E 3 -g 4 target.bed \| teststrandbias.R \| var2vcf_valid.pl -N output -f 0.01 > output.vcf | (19) |
| VarScan2 | A robust heuristic method with statistical test, based on pre-defined thresholds | SNVs and INDELs | v2.4.2 | samtools mpileup -f ref.fa -d 1000 -l target.bed -E -q 20 -Q 20 input.bam > output.pileup;<br>java -jar VarScan.v2.4.2.jar mpileup2snp output.pileup --min-coverage 5 --min-reads2 2 --min-avg-qual 20 --min-var-freq 0.002 --output-vcf 1 > output.SNP.vcf;<br>java -jar VarScan.v2.4.2.jar mpileup2indel output.pileup --min-coverage 5 --min-reads2 2 --min-avg-qual 20 --min-var-freq 0.002 --output-vcf 1 > output.INDEL.vcf | (20) |
| Platypus | Local de novo assembly, Bayesian statistical framework | SNVs and INDELs | v0.5.2 | python Platypus.py callVariants --bamFiles=input.bam --refFile=ref.fa --ploidy=2 --nCPU=n_threads --maxVariants=15 --minBaseQual=17 --output=output.vcf --regions=target.bed | (21) |
| Freebayes | Haplotype-based, Bayesian statistic | SNVs, MNPs and short INDELs | v1.3.1 | freebayes input.bam -v output.vcf -f ref.fa -t target.bed -C 2 -3 20 -P 0.0001 -q 17 -W 1,3 -S 4 -M 3 -B 25 -E 3 | (22) |
| GATK Mutect2 | Local assembly of haplotypes, Bayesian genotyping model | SNVs and short INDELs | v3.7 | java -jar GenomeAnalysisTK.jar -T MuTect2 -R ref.fa -L target.bed -I:tumor input.bam -dt NONE -o output.vcf | (23) |

| | | | | | |
|---|---|---|---|---|---|
| GATK HC | Local assembly of haplotypes, Bayesian statistic | SNVs and short INDELs | v3.7 | java -jar GenomeAnalysisTK.jar -T HaplotypeCaller -R ref.fa -L target.bed -I input.bam -ploidy 2 -stand_call_conf 0 -o output.vcf | (24,25) |
| GATK UG | Bayesian genotype likelihood model | SNVs and short INDELs | v3.3 | java -jar GenomeAnalysisTK.jar -T UnifiedGenotyper -R ref.fa -L target.bed -glm BOTH -mbq 17 -ploidy 2 -stand_call_conf 0 -stand_emit_conf 0 -I input.bam -o output.vcf | (24,25) |
| Strelka2 | Estimation of INDEL error parameters based on a Bayesian mixture model, tiered haplotype model | SNVs and short INDELs | v2.8.3 | configureStrelkaGermlineWorkflow.py --bam input.bam --referenceFasta ref.fa --callMemMb=1024 --exome --disableSequenceErrorEstimation --runDir output; runWorkflow.py -m local -j 10 | (26) |
| mpileup | Bayesian inference, available in bcftools | SNVs and short INDELs | v1.9 | samtools mpileup -B -Q 1 -C 0 -d 2000 --per-sample-mF -l target.bed --output-tags DP,AD -f ref.fa --BCF input.bam \| /path_to/bcftools call --multiallelic-caller --variants-only -O z > output.vcf.gz | (27) |
| VarTracker | Read count-based, utilize bcftools mpileup | SNVs and short INDELs | NA | bcftools mpileup -C 0 -d 1000 -r $target --output-type v -f ref.fa --output output.vcf input.bam; $target is a target region in the format of chr:start-end | This study |

GATK UG, UnifiedGenotyper; GATK HC, HaplotypeCaller; SV, structural variants

**Supplementary Table S4**. Features used to build ML variant classifier

| Category | Feature | Short name | Description |
|---|---|---|---|
| Variant quality | 1 | FS* | Phred-scaled p-value from Fisher's exact test of strand bias |
| | 2 | GC | GC content within $\pm 20\text{bp}$ of a given variant position |
| | 3 | HRun | Largest contiguous homopolymer run of variant allele in either direction (numeric) |
| | 4 | MQ | Average mapping quality of variant-carrying reads |
| | 5 | MQ0* | Number of variant-carrying reads with mapping quality of 0 |
| | 6 | NBase | Percent of unknown (N) bases at the variant position in the pileup |
| | 7 | STR | Variant is part of a short tandem repeat (0-no, 1-yes) |
| | 8 | DP | Read depth at a given variant position |
| | 9 | DPref | Read depth supporting the reference allele |
| | 10 | DPalt | Read depth supporting the alternative allele |
| | 11 | VAF | Variant allele frequency |
| Variant caller | 12 | Mutect | Mutect2 calling status (0-no, 1-yes) |
| | 13 | VarDict | VarDict calling status (0-no, 1-yes) |
| | 14 | VarTracker | VarTracker calling status (0-no, 1-yes) |
| | 15 | MetaCaller | MetaCaller calling status (1-by one caller, 2-by two callers, 3-by all three callers) |
| Mutation signature | 16 | MutSig | Mutational signature (0-C->T, 1-others) |
| Genomic context | 17 | Mscore | Mappability score of the variant position (range: 0-1) |
| | 18 | Repeat | Repeat masked region (0-no, 1-yes) |
| | 19 | LCR | Low complexity region (0-no, 1-yes) |
| | 20 | CpG | CpG island (0-no, 1-yes) |
| | 21 | SD | Segmental duplication region (0-no, 1-yes) |
| Population data | 22 | gnomAD.MAF | Overall MAF in gnomAD |
| | 23 | 1000G.MAF | Overall MAF in 1000 Genomes Project |
| | 24 | TOPMed.MAF | Overall MAF in TOPMed (NHLBI Trans-Omics for Precision Medicine) |
| | 25 | rsID | Reference SNP in dbSNP (0-no match, 1-with match) |
| | 26 | COSMIC.FREQ | Number of samples carrying the variant in the COSMIC database |

Categorical variables include feature 7, 12-16, 18-21, and 25.

*FS and MQ0 are not applicable to INDELs
MAF, minor allele frequency
Mappability score, only keep positions with mappability score ≤0.5 (i.e., subsequences that occur at least twice in the genome)
Mappability score, http://hgdownload.cse.ucsc.edu/gbdb/hg19/bbi/wgEncodeCrgMapabilityAlign100mer.bw
Repeat, https://genome.ucsc.edu/cgi-bin/hgTables (specify assembly:hg19, group:Repeats, track:RepeatMasker)
LCR, https://github.com/lh3/varcmp/raw/master/scripts/LCR-hs37d5.bed.gz
CpG, https://genome.ucsc.edu/cgi-bin/hgTables (specify assembly:hg19, group:Regulation, track:CpG Islands)
SD, https://eeelegacy.gs.washington.edu/humanparalogy/build37/data/GRCh37GenomicSuperDup.tab
gnomAD, http://gnomad.broadinstitute.org/downloads (select gnomAD v2.1.1, Build=GRCH37)
1000G, ftp.1000genomes.ebi.ac.uk/vol1/ftp/release/20130502 (select *.20130502.genotypes.vcf.gz files)
TOPMed, https://bravo.sph.umich.edu/freeze3a/hg19/
rsID, https://ftp.ncbi.nih.gov/snp/archive/ (select b152/)
COSMIC, https://cancer.sanger.ac.uk/cosmic/download (select GRCh37, v95)

**Supplementary Table S5**. Machine learning tools assessed for CHIP prediction

```
library(tidyverse)   # data manipulation
library(mlr)           # ML package
library(xgboost)    # ML package
library(caret)       # generate confusion matrix
library(knitr)        # make pretty tables

#### Rpart and ksvm

## read feature data
feature_df=read.table(file="feature.txt", sep="\t", header=T)

## build class
var.task = makeClassifTask(data=feature_df,target="CLASS_LABEL")

## get training and test dataset
set.seed(1000)
n = getTaskSize(var.task)
train.set = sample(n, size = round(0.8 * n))
test.set = setdiff(seq_len(n), train.set)

## create learner object
# for rpart
lrn1 = makeLearner("classif.rpart", predict.type = "prob
# for ksvm
lrn1 = makeLearner("classif.ksvm", par.vals = list(kernel = "vanilladot"))

## train model and make prediction
mod1 = mlr::train(lrn1, var.task, subset = train.set)
pred1 = predict(mod1, task = var.task, subset = test.set)

## generate confusion matrix
cmat=confusionMatrix(as.factor(pred1$data$response), as.factor(pred1$data$truth))
```

```
#### Random forest at default parameters

## build class, and get training and test dataset as in Rpart and ksvm

## create learner object
lrn1 = makeLearner("classif.randomForest", predict.type = "prob")

## train model and make prediction at default parameters
mod1 = mlr::train(lrn1, var.task, subset = train.set)
pred1 = predict(mod1, task = var.task, subset = test.set)

## generate confusion matrix
cmat=confusionMatrix(as.factor(pred1$data$response), as.factor(pred1$data$truth))

#### Random forest with hyperparameter tuning

## build class, get training and test dataset, and create learner object as above

## grid search to find hyperparameters
rf_param <- makeParamSet(
makeIntegerParam("ntree",lower = 50, upper = 500),
makeIntegerParam("mtry", lower = 3, upper = 10),
```

```
makeIntegerParam("nodesize", lower = 10, upper = 50)
)

## search for 50 iterations
rancontrol <- makeTuneControlRandom(maxit = 50L)

## set 5-fold cross validation
set_cv <- makeResampleDesc("CV",iters = 5L)

## hyperarameter tuning
rf_tune <- tuneParams(learner = rf, resampling = set_cv, task = trainTask, par.set = rf_param, control =
rancontrol, measures = acc)

## use hyperparameters for modeling
rf.tree <- setHyperPars(rf, par.vals = rf_tune$x)

## train model and make prediction
rforest <- mlr::train(rf.tree, trainTask)
rfmodel <- predict(rforest, testTask)

## get confusion matrix
cmat=confusionMatrix(rfmodel$data$response, rfmodel$data$truth)
```

**#### XGBoost at default parameters**

```
## read feature data
feature_df=read.table(file="feature.txt", sep="\t", header=T)

## get training and test dataset
n = nrow(feature_df)
train.index = sample(n,floor(0.8*n))

train.data = as.matrix(feature_df[train.index,])
mode(train.data)="numeric"
test.data = as.matrix(feature_df[-train.index,])
mode(test.data)="numeric"

train.label = label[train.index]
test.label = label[-train.index]

traintask <- makeClassifTask (data = train.data, target = "CLASS_LABEL")
testtask <- makeClassifTask (data = test.data, target = "CLASS_LABEL")

## define class
# Create an xgboost learner and output labels (as opposed to probabilities)
num_class = length(levels(class_label))
set.seed(123)

xgb_learner <- makeLearner(
  "classif.xgboost",
  predict.type = "prob",
  par.vals = list(
    objective = "multi:softprob",
    eval_metric = "mlogloss",
    nrounds = 298,
    nthread=14
  )
)
```

```
## train model
m0.xgb_model <- mlr::train(xgb_learner, traintask)

## predict outcomes with the test data
m0.xgb.pred = predict(m0.xgb_model, testtask,reshape=T)
m0.xgb.pred = as.data.frame(m0.xgb.pred)

## get confusion matrix
cmat=confusionMatrix(as.factor(m0.xgb.pred$response), as.factor(m0.xgb.pred$truth))

#### XGBoost with hyperparameter tuning
## get training and test dataset, and define class as above

## parameter setup
xgb_params <- makeParamSet(
  # The number of trees in the model (each one built sequentially)
  makeIntegerParam("nrounds", lower = 100, upper = 500),
  # number of splits in each tree
  makeIntegerParam("max_depth", lower = 3, upper = 15),
  makeNumericParam("min_child_weight",lower = 1L,upper = 15L),
  makeNumericParam("subsample",lower = 0.5,upper = 1),
  makeNumericParam("colsample_bytree",lower = 0.5,upper = 1),
  # "shrinkage" - prevents overfitting
  makeNumericParam("eta", lower = 0.1, upper = 0.5),
  # L2 regularization - prevents overfitting
  makeNumericParam("lambda", lower = -1, upper = 0, trafo = function(x) 10^x)
)

## control
control <- makeTuneControlRandom(maxit = 100)
resample_desc <- makeResampleDesc("CV", iters = 5)

## hyperparameter tuning based prediction
tuned_params <- tuneParams(
  learner = xgb_learner,
  task = traintask,
  resampling = resample_desc,
  par.set = xgb_params,
  control = control
)

# create a new model
xgb_tuned_learner <- setHyperPars(
  learner = xgb_learner,
  par.vals = tuned_params$x
)

# re-train parameters using tuned hyperparameters
m1.xgb_model <- mlr::train(xgb_tuned_learner, traintask)

# predict outcomes with the test data
m1.xgb.pred = predict(m1.xgb_model, testtask,reshape=T)
m1.xgb.pred = as.data.frame(m1.xgb.pred)

## get confusion matrix
cmat=confusionMatrix(as.factor(m1.xgb.pred$response), as.factor(m1.xgb.pred$truth))
```

```
#### Build a neural network model with TensorFlow

import pandas as pd
import torch
import numpy as np
import scipy
import tensorflow as tf
import matplotlib.pyplot as plt
import sklearn.metrics
import sklearn.model_selection

## read in feature
filename = "feature.txt"
df = pd.read_csv(filename, delimiter = "\t")
print(df.shape)
df.head(10)

df = df.fillna(0)
print(df.shape)

features = df.to_numpy()[:,1:].astype(np.float64)
labels = df.to_numpy()[:,0]

# quantify labels
label_dict = {"ARTIFACTS": 0, "CHIP": 1, "GERMLINE": 2}
labels = np.array([label_dict[label] for label in labels]).astype(np.int8)

print(features, labels)
print(features.shape, labels.shape)

## get training and test dataset
X_train, X_test, Y_train, Y_test = sklearn.model_selection.train_test_split(features, labels,
train_size = 0.8, random_state = 42)

## normalization
scaler = sklearn.preprocessing.StandardScaler().fit(X_train)
X_train = scaler.transform(X_train)
X_test = scaler.transform(X_test)

## build model
model = tf.keras.models.Sequential([
  tf.keras.layers.Dense(2, activation='relu'),
  tf.keras.layers.Dense(3)
])

## train model
loss_fn = tf.keras.losses.SparseCategoricalCrossentropy(from_logits=True)

model.compile(optimizer='adam',
              loss=loss_fn,
              metrics=['accuracy'])

model.fit(X_train, Y_train, validation_data = (X_test, Y_test), epochs=5)

## predict and get confusion matrix
from sklearn.metrics import confusion_matrix, recall_score, precision_score

Y_pred = np.argmax(model.predict(X_test), axis = -1)
conf_matrix = confusion_matrix(Y_test, Y_pred)
```

```
print(conf_matrix.T )

# prediction is on y-axis, reference is on x-axis
print(sum(Y_test == 2))

plt.imshow(conf_matrix, cmap = plt.cm.Greens)
plt.xlabel("Predicted")
plt.ylabel("True")
```

**Supplementary Table S6.** Performance metrics of neural network in CHIP prediction

| Type | Recall (%) | Precision (%) | Accuracy (%) |
|---|---|---|---|
| SNV | 64.63 | 86.78 | 82.05 |
| INDEL | 90.26 | 91.07 | 87.92 |

Recall, precision and accuracy are estimated using formula (2), (1) and (5), respectively. Neural network was tested separately on SNV and INDEL, using WGS data from batch 2 simulation with CHIP-specific VAFs (Supplementary Table S2).

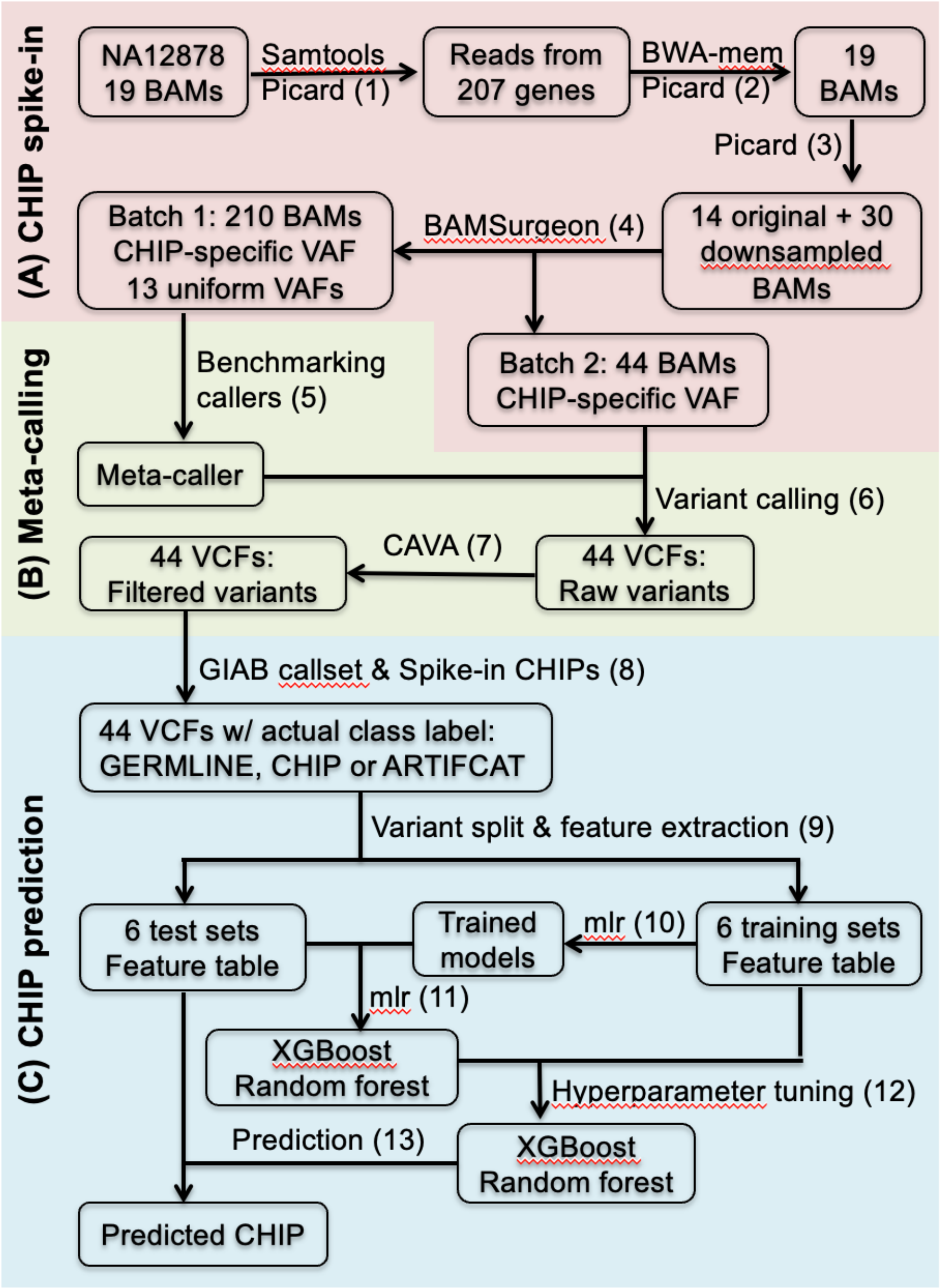

**Supplementary Figure S1**. Flowchart for developing the meta-caller and variant classifier. (**A**) CHIP spike-in (step 1-4). NA12878 WES and WGS alignments within the 202 leukemia-associated genes were extracted, remapped to the hg19 genome reference, and then downsampled to a minimum coverage of 20X (Supplementary Table S2). Of the resulting 44 BAMs, 21 were used as inputs in batch 1 simulation, as shown in Supplementary Table S2. The 2,367 known CHIP were added as spike-in at CHIP-specific VAFs, or at each of 13 uniform VAFs between 0.5% and 30%. Batch 2 simulation was performed on all 44 BAMs using CHIP-specific VAFs.

(**B**) Variant detection using meta-caller (step 5-7). Batch 1 simulated data were used to benchmark ten open source tools plus VarTracker. The meta-caller was built by combining VarTracker with Mutect2 and Vardict, the two top-performing tools. Variants were called from batch 2 simulation data and used to develop the variant classifier. (**C**) CHIP prediction (step 8-13). Raw variants with functional effects (via CAVA annotation) were each assigned an actual class label of GERMLINE, CHIP or ARTIFCAT; the associated features were extracted from VCF (Supplementary Table S4). The annotated variants were then used as the training sets (80% of the data) to build machine learning-based prediction models in the feature space, separately for variant type (SNV and INDEL) and sequencing type (WES, WGS, and both combined). XGBoost performed well (with the highest F1-score) on predicting CHIP in the test sets (20% of the data), followed by Random forest. Both were selected for hyperparameter tuning, and used to predict CHIP in the test sets. Numbers in the parenthesis indicate analysis steps in order. SNV, single nucleotide variant.

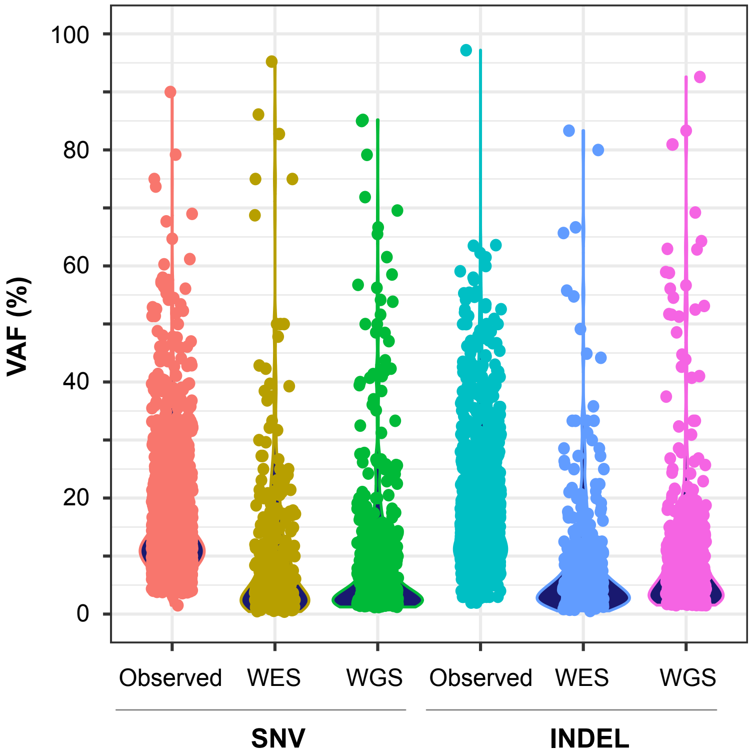

**Supplementary Figure S2**. VAF distribution for known and simulated CHIP mutations. For simulated data with CHIP-specific VAFs, VAFs were estimated based on read pileup at sites carrying pre-inserted mutations. WES, WES_NA12878_01 simulated at 50X; WGS, WGS_NA12878_01 simulated at 50X; Observed, VAFs from the 2,367 known CHIP (1,331 SNVs and 1,036 INDELs). In both simulated WES and WGS, the spike-in CHIP had reduced VAFs compared to the known CHIP.

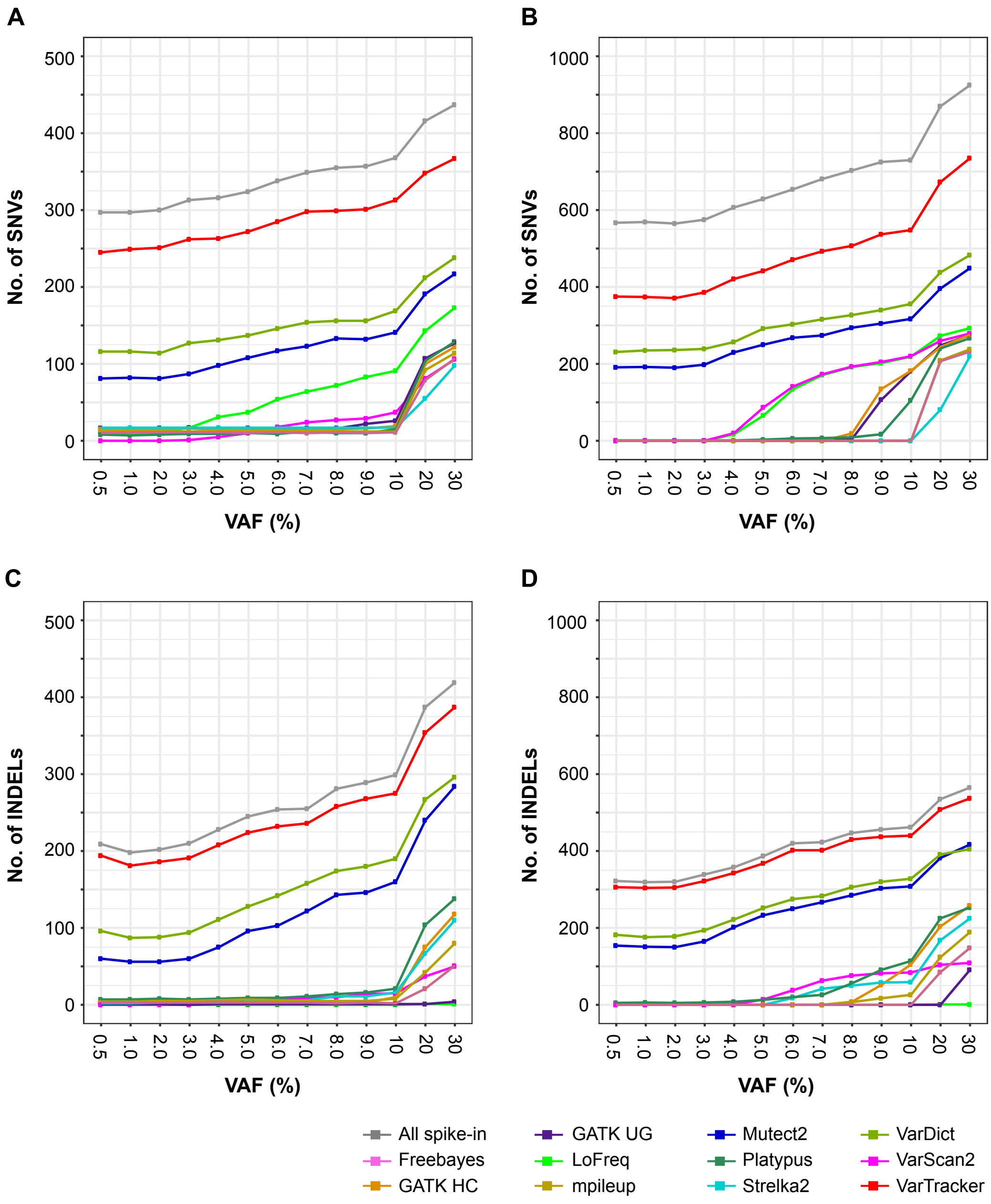

**Supplementary Figure S3**. Recovery of simulated CHIP varies by callers over different VAFs. (**A,C**) SNVs and INDELs simulated at 100X in NA12878_01 WES data. (**B,D**) simulated at 100X in NA12878_01 WGS data. Data are from batch 1 simulation. BAMSurgeon was used to spike-in known SNVs/INDELs at each of the 13 uniform VAFs between 0.5% and 30% (X-axis). Y-axis shows the number of total spike-in (gray line) and the number of spike-in that were recovered by individual callers (color lines). VarTracker, followed by VarDict and GATK Mutect2, identified the largest number of spiked-in, particularly at low VAFs. See Supplementary Table S2 for details about the two NA12878 samples. GATK UG, UnifiedGenotyper; GATK HC, HaplotypeCaller.

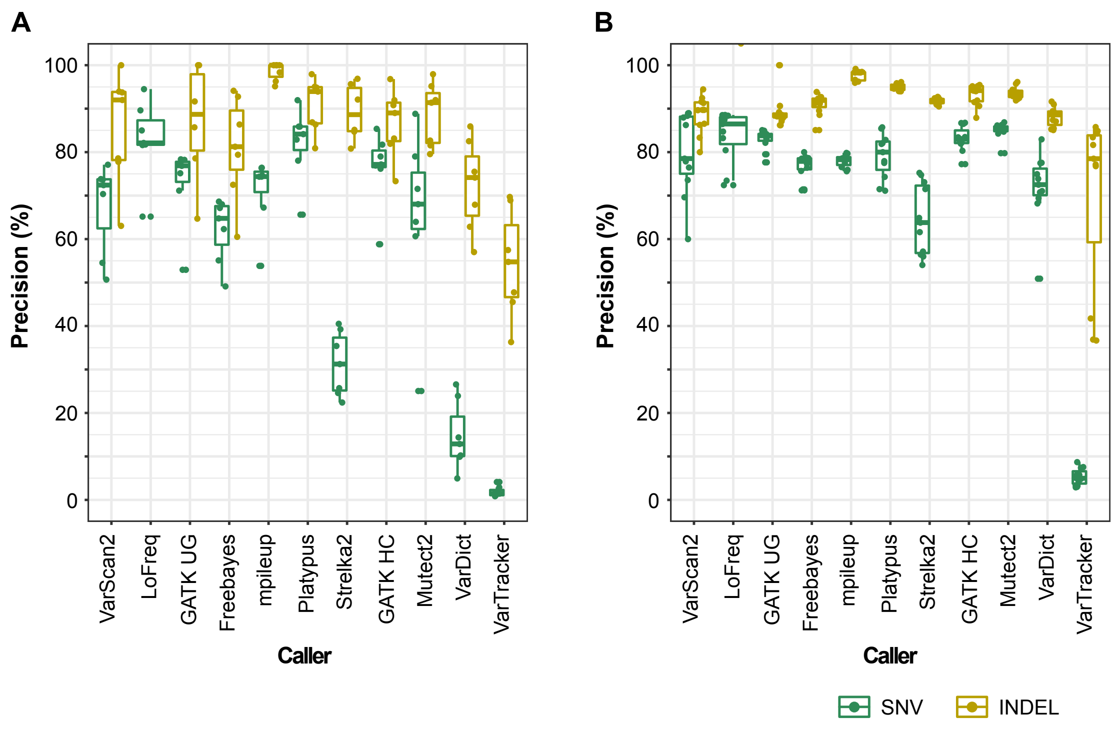

**Supplementary Figure S4**. Precision of 11 tools benchmarked on simulated data. (**A**) Simulated SNVs and INDELs in WES data. (**B**) Simulated SNVs and INDELs in WGS data. Each plot in (A) and (B) used precision estimated from 7 WES and 11 WGS data, respectively, split into SNV and INDEL. All data are from batch 1 simulation (Supplementary Table S2) that used CHIP-specific VAFs, excluding those with >100X coverage. GATK UG, UnifiedGenotyper; GATK HC, HaplotypeCaller.

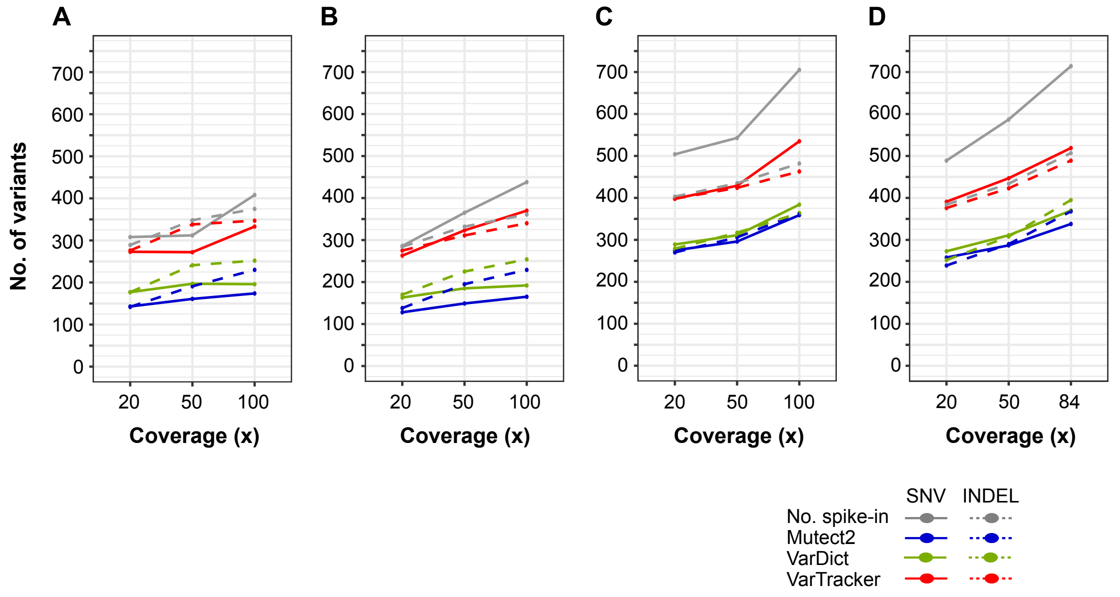

**Supplementary Figure S5**. Recovery of CHIP simulated at different coverage. (**A**) NA12878_01 WES. (**B**) NA12878_02 WES. (**C**) NA12878_01 WGS. (**D**) NA12878_02 WGS. Y-axis shows the number of SNVs and INDELs that were spiked-in (gray lines) and recovered by the three callers (color lines). Overall, the numbers increase following the coverage increase for both WES and WGS. Data are from batch 1 simulation using CHIP-specific VAFs. See Supplementary Table S2 about the WES and WGS data used on simulation.

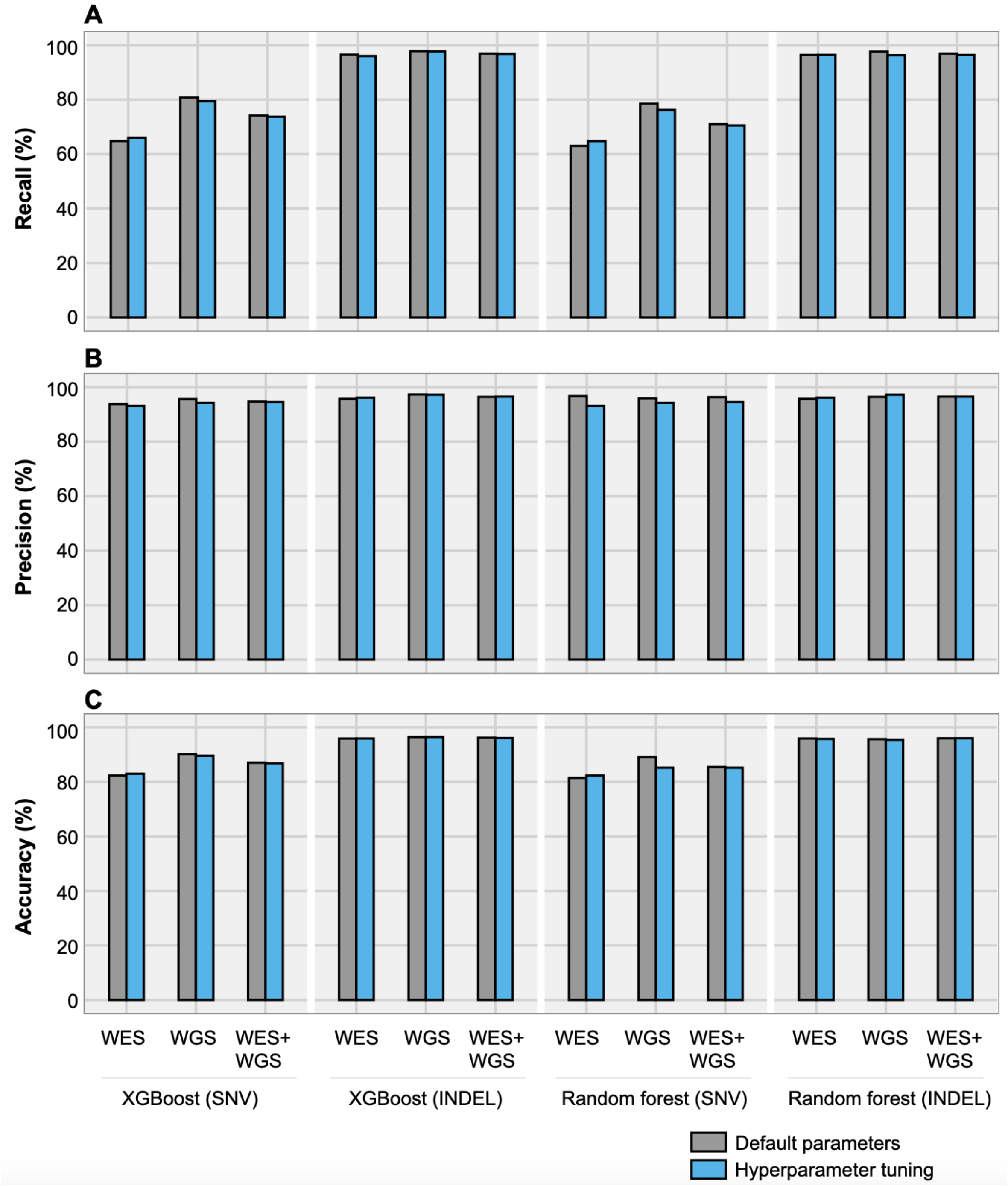

**Supplementary Figure S6.** Performance metrics before and after hyperparameter tuning. (**A**) Recall rate of CHIP prediction based on XGBoost and random forest. (**B**) Precision. (**C**) Accuracy. Recall, precision and accuracy were estimated using formula (2), (1) and (5) respectively. Data were from batch 2 simulation that used CHIP-specific VAFs. Raw variants were called from BAMs by meta-caller, annotated, filtered, and assigned with actual labels of CHIP, GERMLINE, or ARTIFACT.

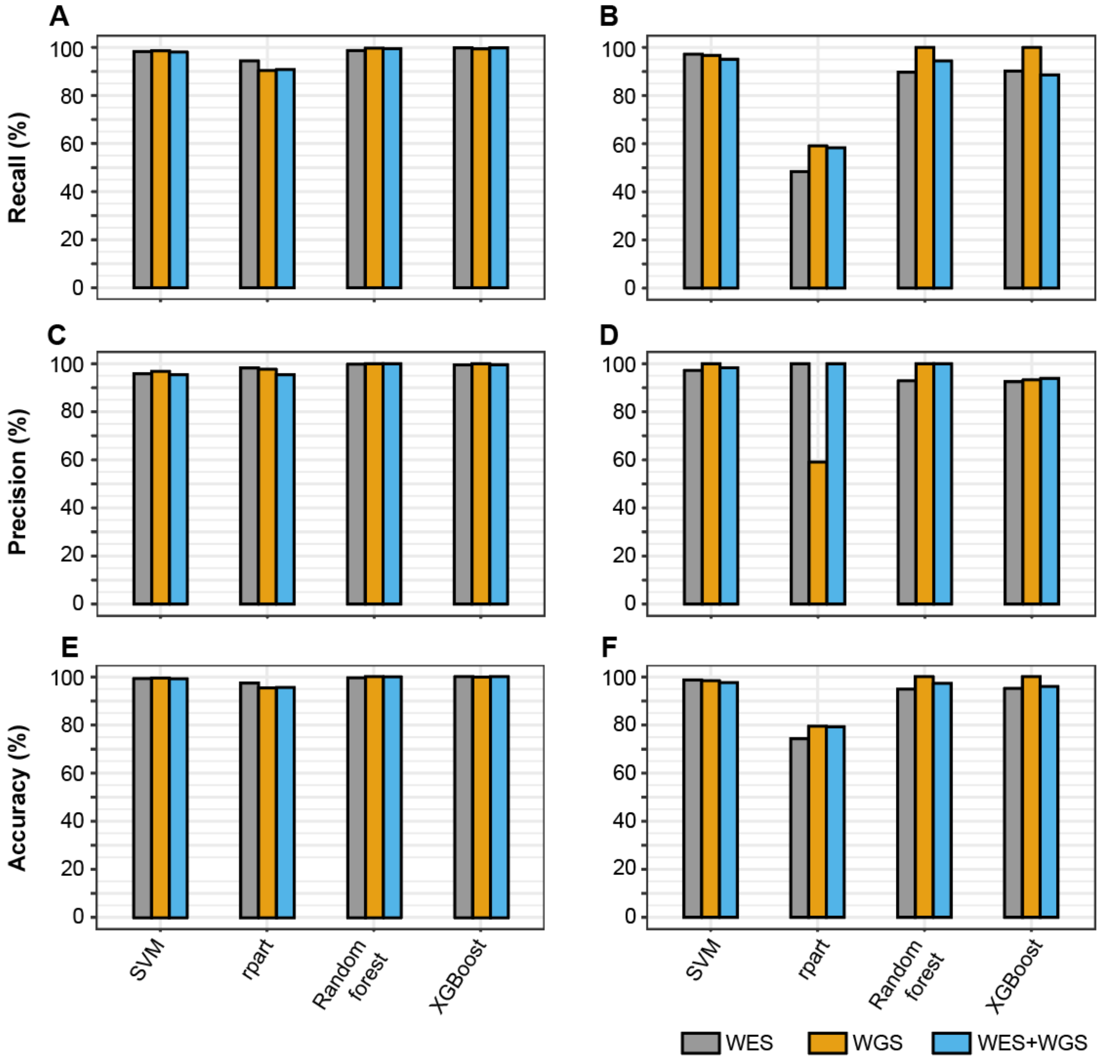

**Supplementary Figure S7**. Performance metrics of machine learning algorithms in predicting germline variants. (**A**) Recall of SNV prediction. (**B**) Recall of INDEL prediction. (**C**) Precision of SNV prediction. (**D**) Precision of INDEL prediction. (**E**) Accuracy of SNV prediction. (**F**) Accuracy of INDEL prediction. The prediction models were applied to the lists of CAVA-annotated variants called from WES (25 BAMs), WGS (19 BAMs), and both (44 BAMs) with the Meta-caller, separated into SNVs and INDELs. Each variant in the input VCF was assigned with an actual class label of “GERMLINE”, “CHIP” or “ARTIFCAT”, described in detail in Supplementary note 1 (“Class labels”). Individual steps in simulation (step 1-4), meta-calling (step 6-7) and variant classification (step 8-13) are illustrated in Supplementary Figure S1. Data are from batch 2 simulation using CHIP-specific VAFs. See Supplementary Table S2 for more information about the WES and WGS data.

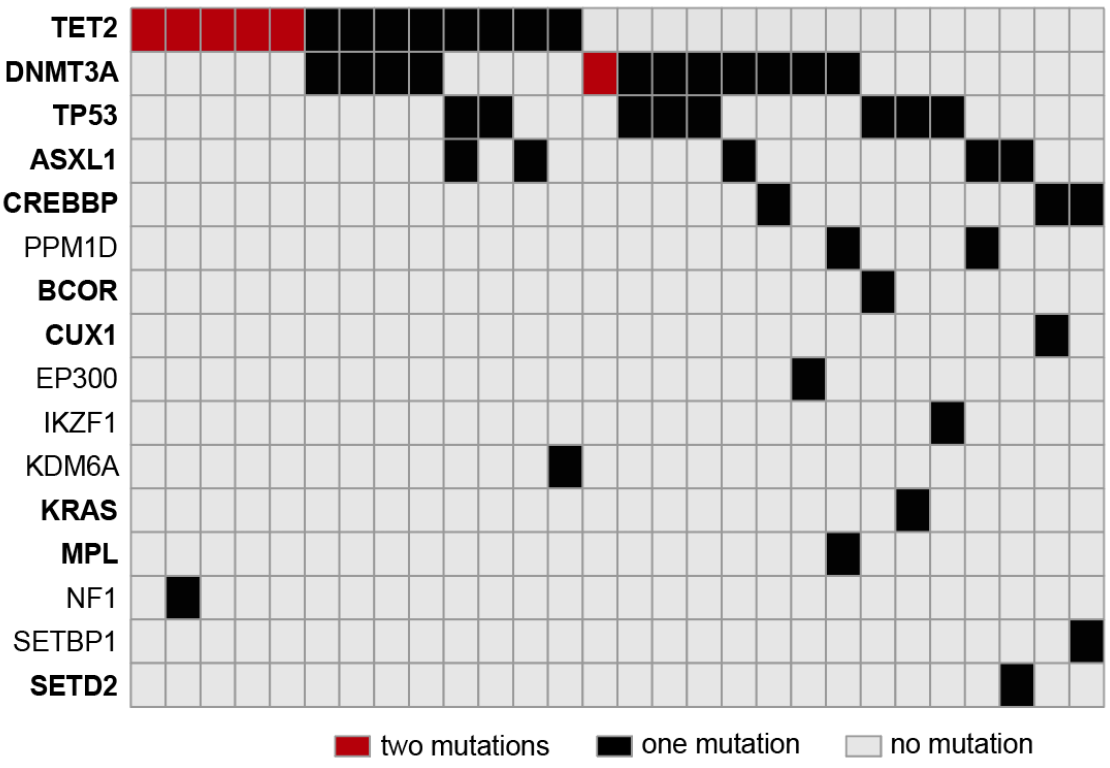

**Supplementary Figure S8**. Plot of individuals with two or more mutations. Columns represent the 28 individuals carrying co-mutation. Ten of the genes (in bold) were found to show co-mutation in a previous study (1).

#### References

1. Jaiswal, S., Fontanillas, P., Flannick, J., Manning, A., Grauman, P.V., Mar, B.G., Lindsley, R.C., Mermel, C.H., Burtt, N., Chavez, A. *et al.* (2014) Age-related clonal hematopoiesis associated with adverse outcomes. *N Engl J Med*, **371**, 2488-2498.
2. Genovese, G., Kahler, A.K., Handsaker, R.E., Lindberg, J., Rose, S.A., Bakhoum, S.F., Chambert, K., Mick, E., Neale, B.M., Fromer, M. *et al.* (2014) Clonal hematopoiesis and blood-cancer risk inferred from blood DNA sequence. *N Engl J Med*, **371**, 2477-2487.
3. Bick, A.G., Weinstock, J.S., Nandakumar, S.K., Fulco, C.P., Bao, E.L., Zekavat, S.M., Szeto, M.D., Liao, X., Leventhal, M.J., Nasser, J. *et al.* (2020) Inherited causes of clonal haematopoiesis in 97,691 whole genomes. *Nature*, **586**, 763-768.
4. Zook, J.M., McDaniel, J., Olson, N.D., Wagner, J., Parikh, H., Heaton, H., Irvine, S.A., Trigg, L., Truty, R., McLean, C.Y. *et al.* (2019) An open resource for accurately benchmarking small variant and reference calls. *Nat Biotechnol*, **37**, 561-566.
5. Kraft, I.L. and Godley, L.A. (2020) Identifying potential germline variants from sequencing hematopoietic malignancies. *Blood*, **136**, 2498-2506.
6. Alexandrov, L.B., Nik-Zainal, S., Wedge, D.C., Aparicio, S.A.J.R., Behjati, S., Biankin, A.V., Bignell, G.R., Bolli, N., Borg, A., Børresen-Dale, A.-L. *et al.* (2013) Signatures of mutational processes in human cancer. *Nature*, **500**, 415-421.
7. Ghoneim, D.H., Myers, J.R., Tuttle, E. and Paciorkowski, A.R. (2014) Comparison of insertion/deletion calling algorithms on human next-generation sequencing data. *BMC Res Notes*, **7**, 864.
8. Bischl, B., Lang, M., Kotthoff, L., Schiffner, J., Richter, J., Studerus, E., Casalicchio, G. and Jones, Z.M. (2016) mlr: Machine Learning in R. *J Mach Learn Res*, **17**, 1-5.
9. Niroula, A., Sekar, A., Murakami, M.A., Trinder, M., Agrawal, M., Wong, W.J., Bick, A.G., Uddin, M.M., Gibson, C.J., Griffin, G.K. *et al.* (2021) Distinction of lymphoid and myeloid clonal hematopoiesis. *Nat Med*, **27**, 1921-1927.
10. Mangaonkar, A.A., Ferrer, A., Pinto, E.V.F., Cousin, M.A., Kuisle, R.J., Gangat, N., Hogan, W.J., Litzow, M.R., McAllister, T.M., Klee, E.W. *et al.* (2019) Clinical Applications and Utility of a Precision Medicine Approach for Patients With Unexplained Cytopenias. *Mayo Clin Proc*, **94**, 1753-1768.
11. Kusne, Y., Lasho, T., Mangaonkar, A., Tefferi, A., Gangat, N., Finke, C., Binder, M., Chia, N. and Patnaik, M.M. (2021) Remarkable stability in clonal hematopoiesis involving leukemia-driver genes in patients without underlying myeloid neoplasms. *Am J Hematol*, **96**, E392-e396.
12. Mouhieddine, T.H., Sperling, A.S., Redd, R., Park, J., Leventhal, M., Gibson, C.J., Manier, S., Nassar, A.H., Capelletti, M., Huynh, D. *et al.* (2020) Clonal hematopoiesis is associated with adverse outcomes in multiple myeloma patients undergoing transplant. *Nat Commun*, **11**, 2996.
13. Ng, P.C. and Henikoff, S. (2003) SIFT: Predicting amino acid changes that affect protein function. *Nucleic Acids Res*, **31**, 3812-3814.

14. Schwarz, J.M., Cooper, D.N., Schuelke, M. and Seelow, D. (2014) MutationTaster2: mutation prediction for the deep-sequencing age. *Nat Methods*, **11**, 361-362.
15. Adzhubei, I., Jordan, D.M. and Sunyaev, S.R. (2013) Predicting functional effect of human missense mutations using PolyPhen-2. *Curr Protoc Hum Genet*, **Chapter 7**, Unit7.20.
16. Xie, M., Lu, C., Wang, J., McLellan, M.D., Johnson, K.J., Wendl, M.C., McMichael, J.F., Schmidt, H.K., Yellapantula, V., Miller, C.A. *et al.* (2014) Age-related mutations associated with clonal hematopoietic expansion and malignancies. *Nat Med*, **20**, 1472-1478.
17. Chen, J., Li, X., Zhong, H., Meng, Y. and Du, H. (2019) Systematic comparison of germline variant calling pipelines cross multiple next-generation sequencers. *Sci Rep*, **9**, 9345.
18. Wilm, A., Aw, P.P., Bertrand, D., Yeo, G.H., Ong, S.H., Wong, C.H., Khor, C.C., Petric, R., Hibberd, M.L. and Nagarajan, N. (2012) LoFreq: a sequence-quality aware, ultra-sensitive variant caller for uncovering cell-population heterogeneity from high-throughput sequencing datasets. *Nucleic Acids Res*, **40**, 11189-11201.
19. Lai, Z., Markovets, A., Ahdesmaki, M., Chapman, B., Hofmann, O., McEwen, R., Johnson, J., Dougherty, B., Barrett, J.C. and Dry, J.R. (2016) VarDict: a novel and versatile variant caller for next-generation sequencing in cancer research. *Nucleic Acids Res*, **44**, e108.
20. Koboldt, D.C., Zhang, Q., Larson, D.E., Shen, D., McLellan, M.D., Lin, L., Miller, C.A., Mardis, E.R., Ding, L. and Wilson, R.K. (2012) VarScan 2: somatic mutation and copy number alteration discovery in cancer by exome sequencing. *Genome Res*, **22**, 568-576.
21. Rimmer, A., Phan, H., Mathieson, I., Iqbal, Z., Twigg, S.R.F., Wilkie, A.O.M., McVean, G., Lunter, G. and Consortium, W.G.S. (2014) Integrating mapping-, assembly- and haplotype-based approaches for calling variants in clinical sequencing applications. *Nat Genet*, **46**, 912-918.
22. Garrison, E. and Marth, G. (2012). http://arxiv.org/abs/1207.3907v2.
23. Cibulskis, K., Lawrence, M.S., Carter, S.L., Sivachenko, A., Jaffe, D., Sougnez, C., Gabriel, S., Meyerson, M., Lander, E.S. and Getz, G. (2013) Sensitive detection of somatic point mutations in impure and heterogeneous cancer samples. *Nat Biotechnol*, **31**, 213-219.
24. Van der Auwera, G.A., Carneiro, M.O., Hartl, C., Poplin, R., Del Angel, G., Levy-Moonshine, A., Jordan, T., Shakir, K., Roazen, D., Thibault, J. *et al.* (2013) From FastQ data to high confidence variant calls: the Genome Analysis Toolkit best practices pipeline. *Curr Protoc Bioinformatics*, **43**, 11 10 11-11 10 33.
25. DePristo, M.A., Banks, E., Poplin, R., Garimella, K.V., Maguire, J.R., Hartl, C., Philippakis, A.A., del Angel, G., Rivas, M.A., Hanna, M. *et al.* (2011) A framework for variation discovery and genotyping using next-generation DNA sequencing data. *Nat Genet*, **43**, 491-498.
26. Kim, S., Scheffler, K., Halpern, A.L., Bekritsky, M.A., Noh, E., Källberg, M., Chen, X., Kim, Y., Beyter, D., Krusche, P. *et al.* (2018) Strelka2: fast and accurate calling of germline and somatic variants. *Nat Methods*, **15**, 591-594.

27. Li, H., Handsaker, B., Wysoker, A., Fennell, T., Ruan, J., Homer, N., Marth, G., Abecasis, G. and Durbin, R. (2009) The Sequence Alignment/Map format and SAMtools. *Bioinformatics*, **25**, 2078-2079.
