## Supplement2 for "Unified somatic calling and machine learning-based classification enhance the discovery of clonal hematopoiesis of indeterminate potential"

**Supplementary note 2**

### Unified somatic calling and machine learning-based classification enhance the discovery of clonal hematopoiesis of indeterminate potential

#### 1 Supplementary Methods for Statistical Filtering of Variants

##### 1.1 Communication Channel Model

This method examines genomic positions independently and therefore in the subsequent we simplify our notation and discussion by focusing on a single genomic position and the question of evaluating the presence or absence of a somatic SNV at this position. The method assumes that at this position alleles other than homozygous reference are deleterious and thus not polymorphically present in the population. Therefore, we further simplify the problem from detecting specific alternate alleles into a binary problem of reference versus alternative alleles.

Therefore, consider a data set composed of $I$ individuals indexed as $i \in \mathcal{I} := \{1, \ldots, I\}$, who each have sequencing reads observing this genomic position indexed by $n \in \mathcal{N} := \{1, \ldots, N^{(i)}\}$. Each read has three components to its observation: the base $b_n^{(i)}$, the probability of sequencing error at this base $q_n^{(i)}$ (computed from the base quality PHRED score provided by the sequencer), and the probability of mapping error $m_n^{(i)}$ (computed from the mapping quality PHRED score provided by the aligner/mpileup). As a simplification of the raw data, we coarse grain the base $b_n^{(i)}$ to be binary with value zero indicating the base matches the reference and one indicating that it does not match the reference and provides support for an alternate allele.

After deduplication, each sequencing read in a bulk DNA-seq run can be assumed to have originated from a distinct cell with a true-but-unknown allelic state $x_n^{(i)}$ which takes value zero when it is a reference allele and one otherwise. This variable is assumed to be drawn from a Bernoulli distribution with probability $p^{(i)}$, representing the true-but-unknown fraction of alleles in the tissue that differ from the reference allele (i.e., the MAF in the $i$-th patient's tissue that is being sequenced).

We use information theory to help formulate our mathematical model for the observed data

$$\mathcal{D} = \{(b_n^{(i)}, q_n^{(i)}, m_n^{(i)}), n \in \mathcal{N}, i \in \mathcal{I}\}$$

with the goal of outlier detection on the samples' MAFs $p^{(i)}$. Namely, we view the observations as being the outputs of the two-stage asymmetric binary communication channel pictured in Figure 1. The transmission probabilities[1] for a given observation $b_n^{(i)}$ are given by a maximum entropy formulation that produces

$$\begin{aligned} a_1 &= 1 - \frac{m_n^{(i)}}{4} \\ b_1 &= \frac{m_n^{(i)}}{4} \\ c_1 &= \frac{3\ m_n^{(i)}}{4} \\ d_1 &= 1 - \frac{3\ m_n^{(i)}}{4} \\ a_2 &= 1 - \frac{q_n^{(i)}}{3} \\ b_2 &= \frac{q_n^{(i)}}{3} \\ c_2 &= q_n^{(i)} \\ d_2 &= 1 - q_n^{(i)}. \end{aligned}$$

For instance, $a_1, b_1, c_1, d_1$ represent the fact that a mapping error has a one-in-four chance of producing an observation that matches the reference allele if we assume equal probability of any base coming from a mapping error at this position. Likewise, $a_2, b_2$ represent the fact that a well-mapped read from an alternate allele has a one-in-three chance of matching the reference allele when a sequencing error takes place. Finally, $c_2, d_2$ represent the fact that a well-mapped read from a reference allele will always produce an alternate allele if a sequencing error takes place.

As there is no specific need to model the intermediate variable $\tilde{x}_n^{(i)}$ from Figure 1, we can directly model the two stage channel as a single stage channel whose transmission probabilities are given by

$$\begin{aligned} P(b_n^{(i)} = 1 | x_n^{(i)} = 1) &= a_1 a_2 + b_1 c_2 \\ P(b_n^{(i)} = 1 | x_n^{(i)} = 0) &= c_1 a_2 + d_1 c_2, \end{aligned}$$

and clearly $P(b_n^{(i)} = 0 | x_n^{(i)} = 1) = 1 - P(b_n^{(i)} = 1 | x_n^{(i)} = 1)$ while $P(b_n^{(i)} = 0 | x_n^{(i)} = 0) = 1 - P(b_n^{(i)} = 1 | x_n^{(i)} = 0)$. Therefore the marginal probability density for a single observation is a Bernoulli with probability

$$\begin{aligned} P(b_n^{(i)} = 1) &= P(b_n^{(i)} = 1 | x_n^{(i)} = 1) P(x_n^{(i)} = 1) + P(b_n^{(i)} = 1 | x_n^{(i)} = 0) P(x_n^{(i)} = 0) \\ &= (a_1 a_2 + b_1 c_2) p^{(i)} + (c_1 a_2 + d_1 c_2)(1 - p^{(i)}). \end{aligned}$$

#### 1.2 Hierarchical Bayesian Inference

In order to perform inference on the unknown MAFs $p^{(i)}$ using the data $\mathcal{D}$, we will make some modeling assumptions regarding the MAFs and the data. First, to simplify our model, we will assume that the mapping qualities and base qualities are deterministic/error-free values. We furthermore assume that $p^{(i)}$ is a random variable so that we can model our final observation as

$$b_n^{(i)} | p^{(i)} \sim \text{Bernoulli}\left( (a_1 a_2 + b_1 c_2) p^{(i)} + (c_1 a_2 + d_1 c_2)(1 - p^{(i)}) \right). \tag{1}$$

However this requires us to put a prior on $p^{(i)}$, which we select as

$$p^{(i)} | \alpha, \beta \sim \text{Beta}(\alpha, \beta)$$

where the hyper parameters $\alpha, \beta$ are also random variables whose prior distribution is given by

$$\begin{aligned} \alpha &\sim \text{InverseGamma}(2, 2) \\ \beta &\sim \text{InverseGamma}(2, 10) \end{aligned}$$

[1]For notational simplicity, we do not include the dependence on $i$ and $n$ for the transmission probabilities $a_1, a_2, b_1, b_2, c_1, c_2, d_1, d_2$.

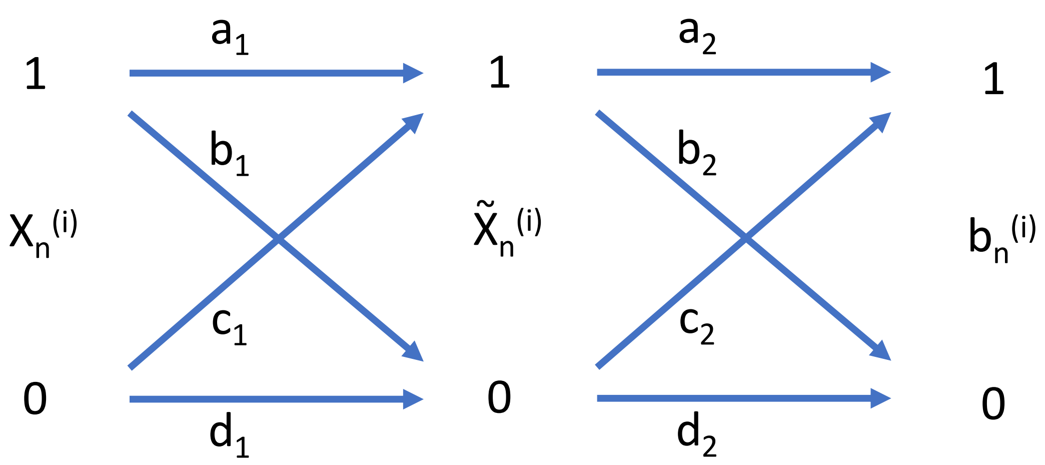


Figure 1: The observed base $b_n^{(i)}$ can be viewed as the output of a two-stage binary communication channel with input $x_n^{(i)}$ being the true-but-unknown allelic state of the cell from which the read is measured. The first stage captures the errors that can arise due to mapping errors, and the second stage represents the errors that can arise due to sequencing errors. Therefore, transmission probabilities $a_1, b_1, c_1, d_1$ are determined by $m_n^{(i)}$ while transmission probabilities $a_2, b_2, c_2, d_2$ are determined by $q_n^{(i)}$.

which have been selected to indicate that for most individuals in the population $p^{(i)}$ is near zero.

For such a complex model, it would be challenging to derive the posterior distribution of $p^{(i)}$ given the observations $b_n^{(i)}$ so rather than proceeding analytically we us the Turing.jl probabilistic programming language to perform Markov Chain Monte Carlo using the No-U-Turn Sampler (NUTS). The major advantage here is that we can jointly sample the vector of MAFs

$$\boldsymbol{p} = \begin{pmatrix} p^{(1)} \\ p^{(2)} \\ \vdots \\ p^{(I)} \end{pmatrix} \tag{2}$$

from the posterior distribution and to therefore estimate the joint distribution of the rank vector

$$\boldsymbol{r} = \begin{pmatrix} \text{rank}_1\, \boldsymbol{p} \\ \text{rank}_2\, \boldsymbol{p} \\ \vdots \\ \text{rank}_I\, \boldsymbol{p} \end{pmatrix} \tag{3}$$

where $\text{rank}_i\, \boldsymbol{p}$ is the descending rank of the $i$-th entry of $\boldsymbol{p}$. Whenever the distribution of $r_i$ concentrates mass near 1, this indicates that the $i$-th individual has unusually large MAF compared to the rest of the population being modeled. Therefore, we use MCMC to compute 95% credible intervals for this rank statistic and deem a sample to be an outlier whenever the lower bound of this credible interval exceeds a given rank threshold $\tau$. Values of $\tau$ closer to 1 will result in fewer outliers being called (lower recall with higher precision) and larger values of $\tau$ wiil result in more outliers being called (higher recall with lower precision). For decently sized populations (i.e., $I > 50$) we suggest $\tau = 0.9I$ as a good heuristic that leads to relatively high precision and recall.
